# The impact of biological sex on alternative splicing

**DOI:** 10.1101/490904

**Authors:** Guy Karlebach, Diogo F.T. Veiga, Anne Deslattes Mays, Christina Chatzipantsiou, Pablo Prieto Barja, Maria Chatzou, Anil K. Kesarwani, Daniel Danis, Georgios Kararigas, Xingmin Aaron Zhang, Joshy George, Robin Steinhaus, Peter Hansen, Dominik Seelow, Julie A McMurry, Melissa A Haendel, Jeremy Yang, Tudor Oprea, Olga Anczukow, Jacques Banchereau, Peter N Robinson

**Affiliations:** The Jackson Laboratory for Genomic Medicine, Farmington, CT 06032, USA; Science and Technology Consulting, LLC, Farmington CT, USA; Lifebit Biotech Ltd., 219 Kensington High Street, London, W86BD, UK; Department of Physiology, Faculty of Medicine, University of Iceland, Reykjavik, Iceland; Berlin Institute of Health, Charitéplatz 1, 10117 Berlin, Germany; Charité-Universitätsmedizin Berlin, Institute of Medical Genetics and Human Genetics, 13353 Berlin, Germany; Linus Pauling institute, Oregon State University, Corvallis OR, USA; Translational Informatics Division, Department of Internal Medicine, The University of New Mexico Health Science Center, Albuquerque, NM 87131, USA; Institute for Systems Genomics, University of Connecticut, Farmington, CT, USA; Department of Genetics and Genome Sciences, UConn Health, Farmington, CT, USA

## Abstract

Over 95% of human genes undergo alternative splicing (AS) in a developmental, tissue-specific, or signal transduction-dependent manner. Here, we present a large-scale survey of sex-biased differential alternative splicing (DAS) across 7027 samples of 39 tissues from 532 individuals (351 males and 181 females) from the Genotype-Tissue Expression project. We detected a total of 1278 statistically significant DAS events affecting 888 different genes and 4417 significant differential gene expression (DGE) events in 3221 genes. Only 267 (29.3%) of the differentially spliced genes were also differentially expressed. Genes that displayed sex-biased DGE or DAS across multiple tissues were enriched in functions related to signaling including histone demethylation. The probability of a gene showing significant differential AS varies by chromosome and is highest for the X chromosome, with differentially spliced X chromosomal genes additionally being more likely to escape X chromosomal inactivation. A small but significant association was found between sex-biased AS and transcripts that undergo physiological nonsense-mediated decay (NMD). We show a significant overlap of differential splicing and genes that display estrogen-induced alternative splicing, that are involved in estrogen response pathway. Further, we show overlap of the involved exons with estrogen-receptor bindings sites. Our results provide a comprehensive survey of sex-biased AS and its characteristics across a large collection of human tissues.

## Introduction

In humans, men and women display important differences in the prevalence, course, and severity of many common diseases, including cardiovascular diseases, autoimmune diseases, and asthma (*1–3*). Despite the importance of sex as a biological variable in development and disease, our understanding of the molecular underpinnings of medically relevant sex differences is still limited. The “sexome” has been defined as the sum of sex-biased effects on gene networks and cell systems. This conceptual framework envisions the functions of cells and tissues as being mediated by a complex and nonlinear network of molecular interactions which increase, decrease, or otherwise modify the activity of nodes (metabolites, chromosomes, proteins and other gene products, etc.) in the network. Sex differences in the network occur because some of the nodes in the network are sensitive to sex-biased factors (*4*).

Biological sex in humans is defined by the sex chromosomes, which are the primary determinants of the sexual differentiation of the gonads and the expression of sex steroid hormones. The primary sex-determining factors are encoded by the sex chromosomes. Y chromosomal genes, such as the testis determining gene *SRY* can only have effects in males. Most X chromosomal genes are subjected to X-chromosomal inactivation in females, a process whereby one of the two copies of an X chromosome is silenced in each female cell through epigenetic modification. Some X chromosomal genes escape X inactivation and thus display constitutively higher expression in female than in male cells (*5*).

Certain genes show sex-biased differential expression during development and in various diseases. Although sex-biased gene expression is a common characteristic of genes encoded on the autosome, chromosome X and Y are enriched for sex-differentially expressed genes (*6*). The mechanisms of sex-biased expression are incompletely understood, but several contributory factors have been described. Gonadal hormones can bind to androgen or estrogen receptors, which in turn can attract cofactors with histone modification activity (*7*). Certain epigenetic modifiers such as the histone demethylase KDM5C show sex-specific expression patterns (*8*), which may provide a partial explanation for the observation that sex-specific DNA methylation is associated with development and disease (*9, 10*).

A number of large-scale studies of sex-specific gene expression have been published including several that investigated data from the Genotype-Tissue Expression (GTEx) project, which currently comprises samples from 54 non-diseased tissue sites assayed by whole genome or exome sequencing and by RNA sequencing (RNA-seq). Data from the GTEx project have been used to study tissue-specific gene expression and regulation (*11–17*). These studies have come to different conclusions, presumably related to different analytical approaches and the fact that earlier studies were restricted to a smaller dataset. One study identified 65 autosomal and 66 X-linked genes with tissue-specific sex differences at a stringent significance threshold (*18*). Another identified 753 genes with tissue specific sex-biased expression predominantly in breast tissue (92%) (*15*). A third found 1559 sex-specific and moderately sex-specific genes (*19*), and the largest study to date found that 37% of all genes exhibit sex-biased expression in at least one tissue (*17*).

In this work, we present a comprehensive investigation of sex-biased alternative splicing (AS). AS, a process by which splice sites are used differentially to create protein diversity, plays an important role in development, disease, and aging (*20–22*). Although some splicing isoforms are produced in the same proportions in all or most cell types, AS is often regulated by developmental or differential cues or in response to external stimuli (*23*). Among the many molecular mechanisms that have been shown to influence AS are Binding of RNA-binding proteins to intronic or exonic regulatory sequences, chromatin-associated factors such as nucleosome density and histone modifications, as well as the splice site sequence itself (*24–27*).

Understanding the effect of both sex-biased differential gene expression and AS is essential for elucidating the molecular correlates of sex differences and may provide a foundation for a better understanding of sex differences in disease. Here, we present a large-scale study of RNA-seq data from the GTEx project to characterize patterns and associations of sex-biased differential splicing.

## Results

### Sex-biased Alternative Splicing and Differential Expression

Here we present an analysis of sex-biased AS based on a comparison of male and female samples from GTEx version 8 (*11*). RNA-seq samples were aligned with hisat2, counts of AS events were determined with rMATS, and sex-biased AS was assessed using linear regression. In parallel, we assessed gene expression in the same samples starting with the transcript-per-million files provided by GTEx and using limma/voom to assess differential gene expression (DGE).

### Biological sex is associated with pervasive differences in gene expression and alternative splicing

As previously reported, DGE between men and women is common (*11–16, 28*). Across the tissues examined in this study, we detected a total of 4417 DGE events in 3221 genes with significantly sex-biased expression at a false-discovery rate (FDR) cutoff of 0.05 and a fold change cutoff of 1.5 (Methods). Clustering analysis revealed a consistent shift in global expression patterns between males and females in some related tissues, such as 9 brain regions, as well as three arterial tissues, two esophageal tissues, and sun-exposed and non-sun-exposed skin (fig. S1).

To analyze differential AS, we focused on five classes of discrete AS events (fig. S2) and defined sex-biased AS based on a statistical model with sex, AS event, and sex:event interaction as covariates. We called AS events sex-biased if the interaction term was significant following multiple testing correction (Methods). There were 1278 statistically significant differential AS (DAS) events over all tissues, affecting 888 different genes.

The overall count of sex-biased DGE and DAS events was strikingly different in different tissue types. In all, there were more instances of DGE than DAS, but breast, thyroid, aorta, and esophagus (muscularis layer) each showed over 100 events. Our analysis revealed between 1 (in pancreas) and 996 (in mammary tissue) significant DAS events per tissue. In addition to mammary tissue, seven other tissues showed over 10 DAS events. As with DGE, DAS was found to be most common in mammary tissue (fig. 1).

**Figure 1:**
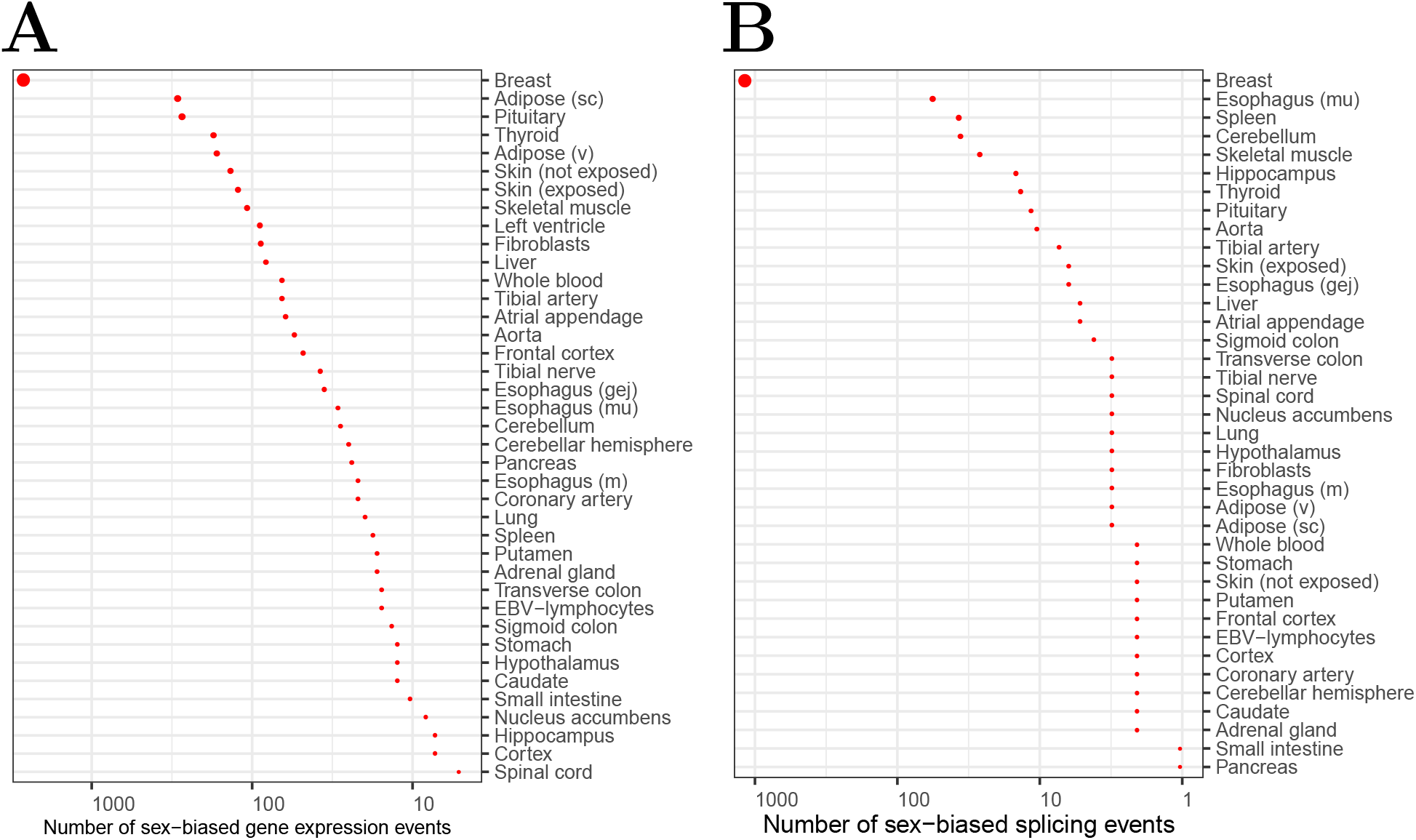
Sex-biased differential expression and splicing. A) Counts of significantly differentially expressed genes per tissue. B) Counts of significantly differentially spliced genes per tissue.

The proportion of alternative skipping event types was skipped exon (SE): 64.2%, retained intron (RI): 9.5%, mutually exclusive exon (MXE): 4.5%, 3’ alternative splice site (A3SS): 13.4%, and 5’ alternative splice site A5SS: 8.4%, with some variation between the chromosomes (fig. 2).

**Figure 2:**
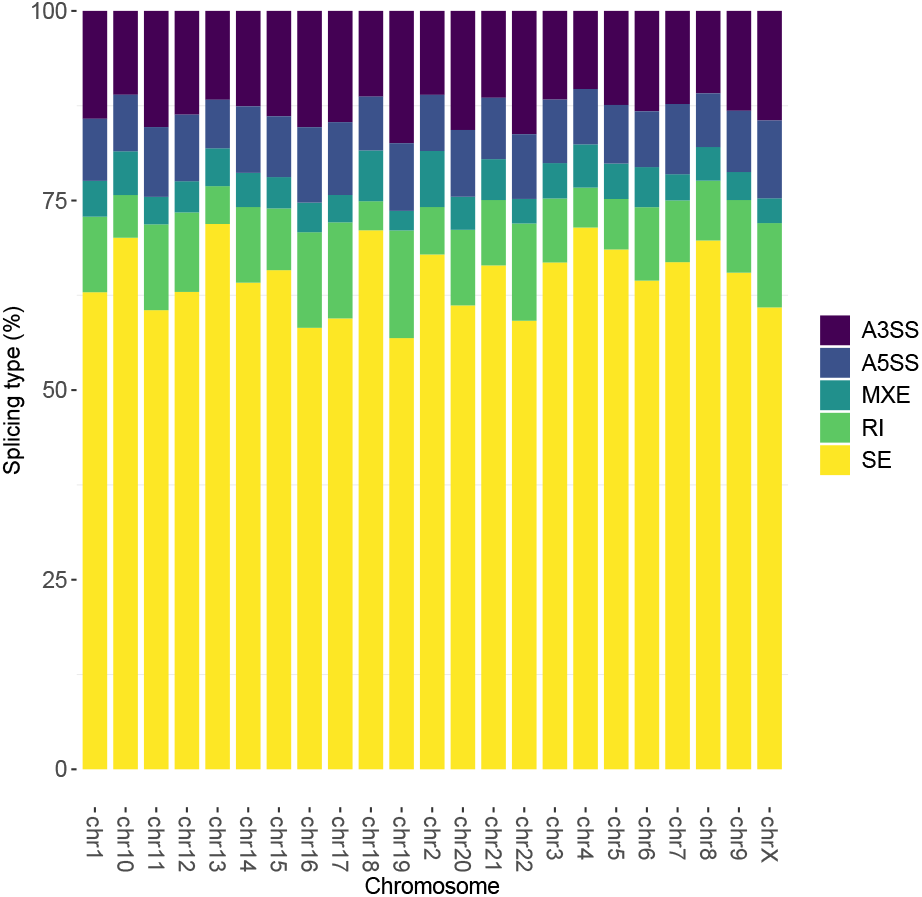
Distribution of different AS events. Skipped-exon (SE) events were the most common followed by 3’ alternative splicing events (A3SS), retained intron events (RI), 5’ alternative splicing events (A5SS), and mutually exclusive exon events (MXE).

### Greater than expected overlap of sex-biased DGE and DAS genes

We assessed the overlap between differential splicing and differential expression by classifying a gene as DGE if it was found to be differentially expressed in at least one tissue, and similarly we classified a gene as DAS if it was found to be differentially spliced in each one tissue. We defined the set of all relevant genes to be those with at least one read in the gene expression dataset (n=24,053). 3221 genes were in the DGE group (13.3%), and 878 (3.6%) were in the DAS group. 2963 genes were DGE but not DAS (92.0% of all DGE genes), and 621 were in the DAS but not the DGE group (70.7% of all DAS genes). This indicates that our definitions of DGE and DAS identified substantially different sets of genes with significant differences between the sexes. However, the overlap between these two sets was higher than one would expect by chance (table 1).

**Table 1:** Differentially spliced and differentially expressed genes. Genes were classified according to whether they were found to be significantly differentially expressed, significantly alternatively spliced, both, or neither. The universe of genes was taken to be all genes with at least one read in the gene expression dataset. 258 (0.62%) of genes were both differentially expressed and differentially spliced, which is more than one would expect by chance (if the two categories were independent, one would expect ~68 genes (0.16%). This observation is statistically significant (Fisher exact test, *p* < 2.2 × 10^−16^).

| DAS | DGE |  |
| --- | --- | --- |
|  | + | - |
| + | 258 | 620 |
| - | 2963 | 20,276 |

It has been reported that differentially expressed sex-biased genes are likely to escape from X chromosome inactivation (*5*). We confirmed this result with our data: 9.8% of all X chromosomal genes escape X inactivation, but 33.9% of differentially expressed X-chromosomal genes do (*p* = 1.70 × 10^−14^, Fisher’s exact test). Additionally, we found that 23.8% of genes displaying sex-biased DAS were associated with escape from X inactivation (*p* = 0.0011) (fig. 3).

**Figure 3:**
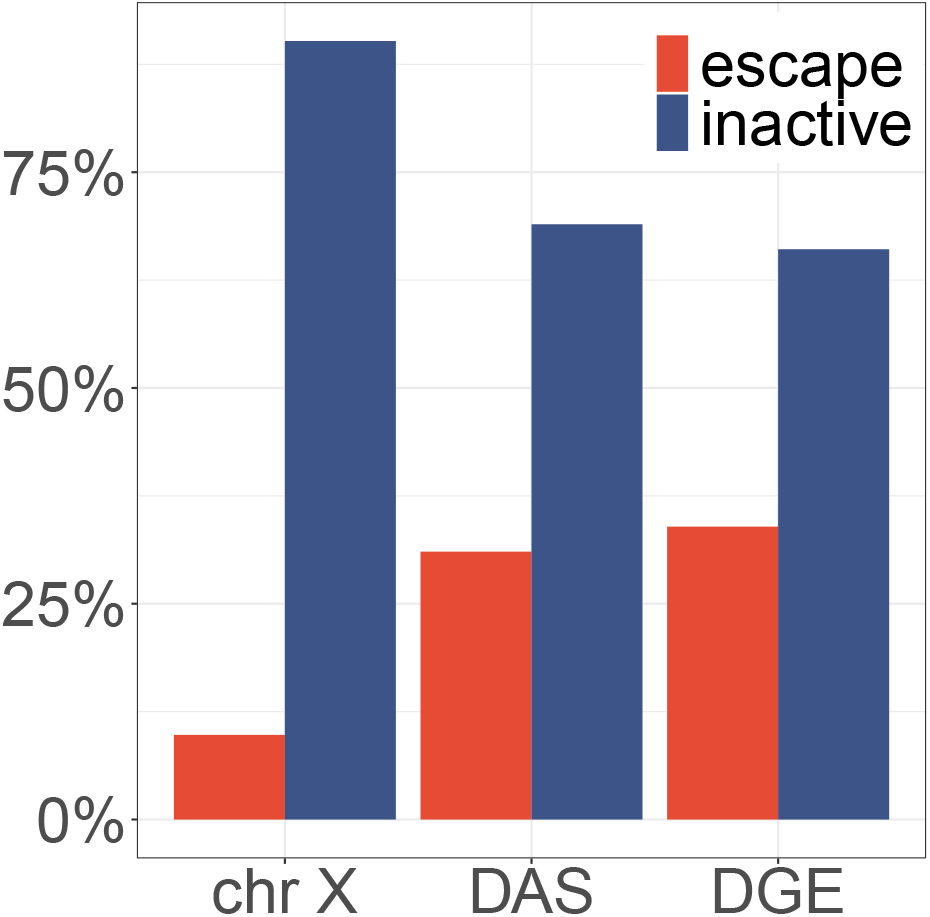
X-chromosomal genes showing differential expression and splicing are more likely to escape X inactivation. 959 X chromosomal genes were investigated by Tukiainen et al. (*5*), 94 of which displayed X chromosomal inactivation. We examined the intersection of DGE and DAS genes. The Y axis shows the percentage of genes that escape X inactivation (escape, red) or genes not shown to display X-inactivation (blue). DAS: X-chromosomal genes showing significantly differential alternative splicing; DGE: X-chromosomal genes showing significantly differential expression.

### Distribution of differential splicing across tissues and chromosomes

The great majority of the DAS events characterized by our study were specific to one of the 39 tissues. 21 events (2.4%) were found in two or more tissues (fig. 4A).

**Figure 4:**
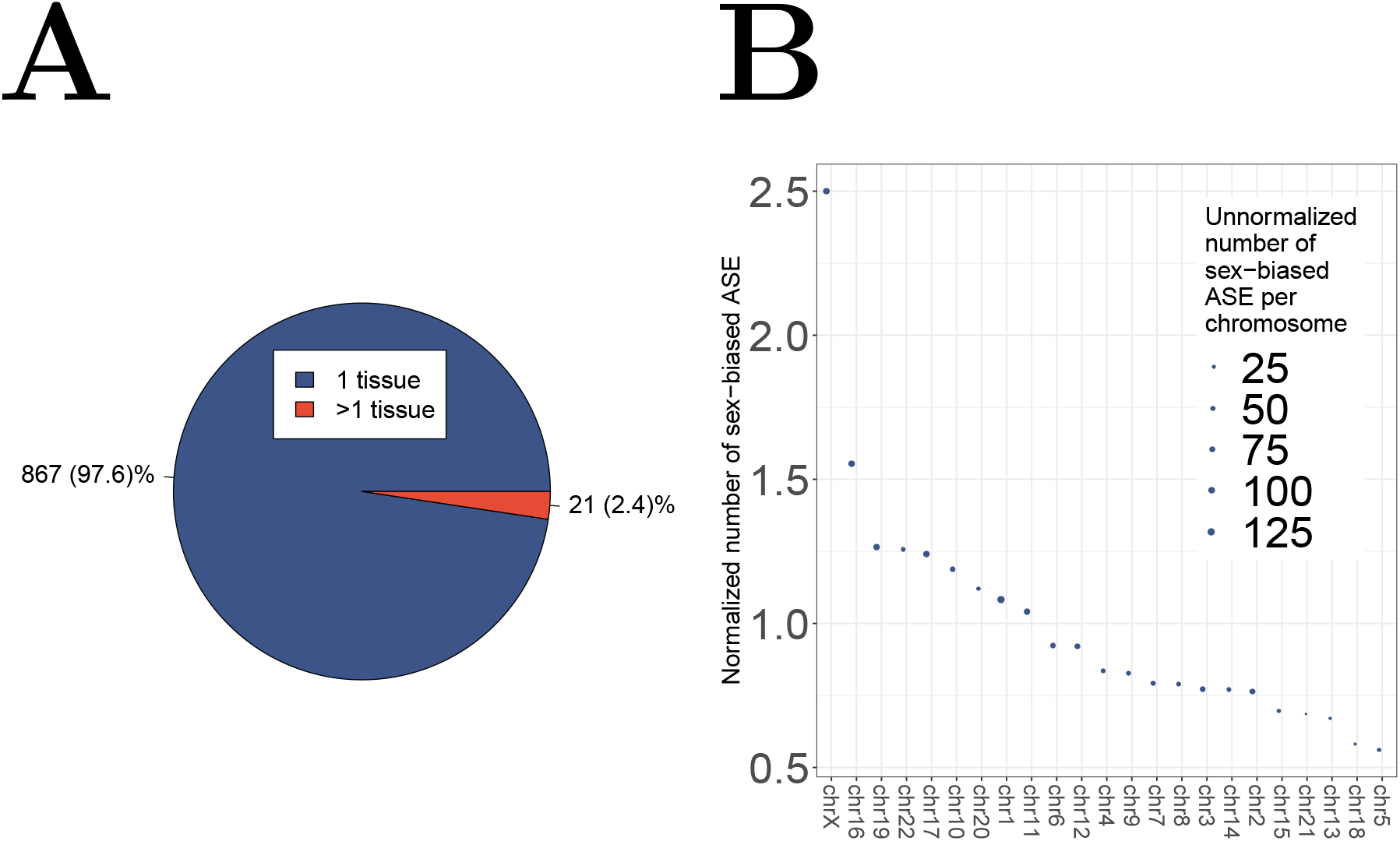
Tissue distribution of DAS. A) The plot shows the distribution of the number of tissues-per-gene for genes with significant differential alternative splicing. B) Sex-biased splicing index (SBSI): the number of significant alternative splicing events per 1000 exons on each chromosome.

Most of the AS events were specific to one tissue. 5 genes showed significant DAS in more than two tissues (*XIST, KDM5C, DDX3X, KDM6A, JPX*). Four of these genes play a role in chromatin remodeling or modification, and all five play roles in regulation of gene expression (table S1). 16 additional genes were significantly differentially spliced in two tissues. Of the top 21 genes, 7 are X chromosomal (table S2). Indeed, the X chromosome showed the highest normalized number of sex-biased AS events per exon. We defined a sex-biased splicing index (SBSI) as the number of statistically significant AS events per 1000 exons, and calculated the SBSI for each chromosome (excluding the Y chromosome). The X chromosome had by far the highest SBSI with 14.6 per 1000 exons showing sex-biased AS events, with most of the remaining chromosomes having an SBSI of between 3 and 5 (fig. 4B).

### Sex-biased alternative splicing affects a diverse set of genes and biological functions

We performed Gene Ontology (GO) (*29*) enrichment analysis to characterize biological functions that are overrepresented among the genes that displayed alternative splicing in each tissue. For comparison, we performed an analogous investigation for DGEs. The proportion of genes annotated to a GO term in each tissue was compared to the population of all 11,270 genes included in the DAS and DGE analysis for which GO annotations were available. 32 of the 39 tissues investigated displayed at least one significant enrichment, with a total of 8808 significant enrichments corresponding to 5229 distinct GO terms (table S4). Fewer terms were found to be significant for the DAS results, which could be related to the lower overall number of genes in the DAS set compared to the DGE set. 8 of the 39 tissues investigated displayed at least one significant enrichment, with a total of 44 significant enrichments corresponding to 42 distinct GO terms. Examples of enriched GO terms include *RNA binding* in the muscularis layer of the esophagus (33/138; 23.9% vs. 10.6% in the population), *response to steroid hormone* in spleen (7/64; 10.9% vs 1.5%), *cadherin binding* in breast (59/1253; 4.7% vs 2.3% in the population), *response to lithium ion* in thyroid (2/17 or 11.8% vs. 0.1%), and *negative regulation of peptidyl-lysine acetylation* in the aorta (2/12;16.7% vs 0.1%).

In many cases, it remains challenging to interpret the biological consequences of alternative splicing because experimental characterization of the biological functions of individual isoforms of genes is lacking. However, some of the alternative splicing events detected affected exons or isoforms with known or likely functional roles. Detailed explanations, visualizations, and references are available in Supplemental Tables S6-S36. A skipped exon (SE) event was found in breast in *ANO1*, which encodes a calcium-activated chloride channel that modulates EGFR- and CAMK-dependent pathways including AKT and MAPK. Females showed a higher degree of skipping, and the event is located in an intracellular loop that plays a role in Ca^2+^ and voltage sensing (fig. S6–S8).

ESRP2 is an epithelial cell-type-specific splicing regulator (*30*). A skipped exon event was identified in breast tissue, with females showing 3 times more inclusion than skipping, a greater difference than males (Figure 5 and S13–S15). Several of the DAS genes are known targets of ESRP2, including *SLK, FGFR2*, and *ESYT2* (*30, 31*) suggesting the possibility that the sex-biased splicing event in ESRP2 could lead to further sex-biased DAS events.

**Figure 5:**
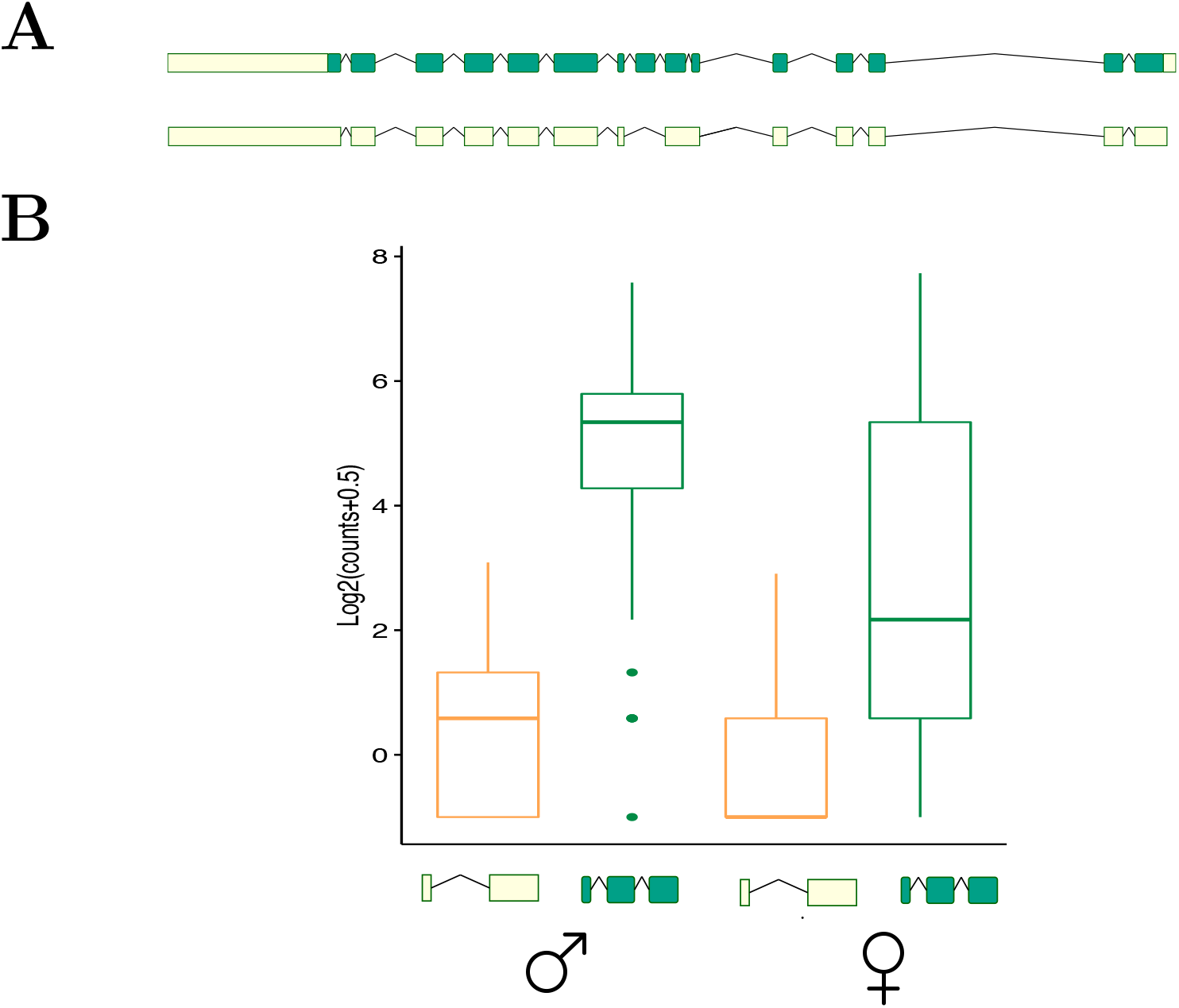
(A) Cartoon showing skip and inclusion isoforms for *ESRP2*. Coding sequences are represented in green, and untranslated regions are shown in yellow. (B) Box plot showing the relative proportions of the skip and inclusion isoforms in males and females in breast tissue (*p* = 6.97 × 10^−1^).

In breast tissue, females displayed greater inclusion of an exon of extended synaptotagmin-2 (*ESYT2*) associated with an increased ability of ESYT2 to elevate cytoplasmic Ca2+ concentration (fig. S16–S18).

Alternative splicing in the gene for fibroblast growth factor receptor 2 (*FGFR2*) generates the IIIb and IIIc isoforms. The IIIb isoforms are expressed exclusively in epithelial cells, while the IIIc isoforms are expressed only in mesenchymal cells. We noted DAS events in breast tissue affecting exons corresponding to the IIIb and IIIc isoforms. Females display a higher degree of inclusion of the exon associated with the IIIb isoform, and a higher degree of skipping of the exon associated with the IIIc isoform. These exons have been considered to be mutually exclusive, but some isoforms exist in Ensembl that contain both exons, which presumably led to rMATS calling two separate SE events (fig.S19–S23). It has previously been reported that mammary epithelial cells express FGFR2-IIIb, which binds FGF7 and FGF10, expressed by surrounding mesenchymal cells (*32*), but sex differences between male and female breast tissue with respect to these isoforms has not been previously investigated.

An SE event in the gene encoded integrin *β*4 (*ITGB4*) was detected in breast tissue (fig. S24– S25). Although no information about the functional specificity of is available, it has been previously determined that distinct isoforms of the most common binding partner of integrin *β*4, integrin *α*6, have different functions (*33*).

The non muscle myosin heavy-chain gene *MYH14* has alternative spliced isoforms that differ by 8 amino acids located in the globular head of the protein. This inclusion is near to the ATP-binding region of MYH14 and increases both actin-activated MgATPase enzymatic activity and motility in translocating actin filaments (*34*). This could suggest a role for MYH14 in the function of myosin containing contractile myoepithelial cells (MEC) involved in lactation (fig. S26–S27).

In breast tissue, females showed a higher level of inclusion of an exon of the transcription factor SLK. The corresponding isoform promotes nuclear localization of SLK (fig. S28–S29). Although the functional consequences thereof are not known, SRSF3-induced inclusion of this exon has also been shown in cancer (*35, 36*).

Several DAS events are can create isoforms that are associated with nonsense-mediated decay. The skipped exons event in *AP3SL* was associated with a shift away from NMD in females (fig. S9–S10). On the other hand, females showed higher levels of NMD-associated isoforms for TBC1D13, a Rab GTPase-activating protein that functions to inhibit insulin-stimulated GLUT4 translocation to the plasma membrane (fig. S30–S31). Females showed higher levels of skipped (non-NMD) isoforms for lysine demethylase 5C (*KDM5C*) in multiple tissues (fig. S32–S36). KDM5C is a ubiquitously expressed X-chromosomal gene that encodes a specific H3K4me3 and H3K4me2 demethylase, and acts as a transcriptional repressor. This suggests the possibility of sex-specific regulation of *KDM5C* by the nonsense-mediated decay pathway (*37*).

Seven distinct alternative splicing events were detected for *DDX3X*. This gene encodes a conserved DEAD-box RNA helicase (fig. S11–S12 and table S7). *XIST* encodes the X-inactive-specific transcript, a long noncoding RNA that plays the key role in inactivating one of the two X chromosomes in females. *XIST* is selectively expressed from and physically coats the future inactive X chromosome in XX female cells (*38*). Experiments in murine embryonic stem cells showed that developmental Xist induction is mediated by enhanced splicing (*39*). XIST displayed significant DAS in 23 tissues, comprising two SE, one A3SS, and one MXE event (fig. S37). As expected, the overall count of reads was substantially higher for females for each of these events. In sigmoid colon, for instance, the mean skip count for a skipped exon event involving exon ENSE00001692247.1 was 0.2 for males (roughly 4.5% of the mean inclusion count of 4.4), and 32.8 for females (8.2% of the mean inclusion count of 400.6) (table S8).

### Sex-biased alternative splicing and Nonsense-Mediated Decay

Nonsense-mediated decay (NMD) is a translation-coupled mechanism that eliminates mRNAs containing premature termination codons (PTCs). NMD can thus serve as a quality control mechanism to prevent the accumulation of abnormal truncated proteins that could be deleterious to the cell (*40*). NMD additionally regulates the abundance of a large number of naturally occurring cellular mRNAs by degrading PTC-containing AS transcripts (*41, 42*). In order to investigate a potential role of such physiological NMD in sex-biased AS, we divided all isoforms of the genes harboring the SE events into isoforms that are predicted to trigger NMD because of the presence of a PTC, and isoforms that do not contain a PTC (which we refer to as non-NMD in the following; Methods). We first tested whether inclusion counts of SE events associated with at least one NMD isoform differed from those of events not associated with any NMD isoform. Of the 786 significantly sex-biased SE events in our dataset, 184 were predicted to be associated with at least one NMD isoform (23.4%), as compared to 28,320 total SE events (significant or not) that passed the minimal count criterion (Methods) out of which 7,538 corresponded to NMD isoforms (26.6%, *p*-value 0.013, Fisher’s exact test). The total number of isoforms associated with NMD that contain a sex-biased event was 238 out of 1506 isoforms containing a sex-biased event (15.8%), as compared to 47,537 transcripts (containing a significant sex-biased event or not) out of which 10,329 are associated with NMD (21.7%). These results show a tendency of sex-biased DAS to result in a different functional isoform in both sexes, although NMD still plays a role in shifting the isoform balance between the sexes.

### Sex-biased alternatively spliced genes in breast tissue significantly overlap with estrogen-induced differentially spliced genes

Estrogen can act throughout the body by several mechanisms including activation of target genes by binding of estrogen receptor (ER). ER*β* binding additionally has been shown to promote alternative splicing of a range of genes (*43*). We therefore hypothesized that estrogen could be associated with sex-biased alternative splicing. To test this hypothesis, we analyzed published data on ER-dependent transcriptional responses in a three-dimensional human breast cell culture model (*44*). We performed isoform-level differential splicing analysis using RSEM (*45*) and HBA-DEALS (*46*). We then computed the overlap between differentially spliced genes for each cell line and genes that contained significant events in breast samples from the GTEx dataset, and combined the hypergeometric p-values using Fisher’s method to obtain a *p*-value of 2.6 × 10^−5^ (table S2). The significant overlap suggests that the sex-biased differential splicing observed in GTEx may also be influenced by estrogen receptor.

### Introns of alternatively-spliced events and estrogen receptor ChIP-Seq peaks significantly overlap

In order to further test the hypothesis that estrogen receptor regulates sex-biased differential splicing, we investigated a set of breast-cancer associated genome-wide ER-binding events that had been assessed by chromatin immunoprecipitation followed by high-throughput sequencing (ChIP-seq) (*47*). We computed the overlap between the flanking introns of sex-biased events in breast tissue and the ESR1 peaks. As a control group, we selected an equal number of events by decreasing p-value, starting from the event with the highest p-value. We then compared the number of base pairs that overlapped with a peak and the number of base pairs that did not between the DAST events and the events in the control group. For the control group these numbers were 675756 and 7084463, respectively, and for the DAST group 853098 and 5119633, which indicates a highly significant overlap between introns of DAST genes and ESR1 binding locations (*p* < 2.2×10^−16^, Fischer’s exact test; table S9). This further supports the notion that estrogen receptor plays a role in sex-biased differential splicing.

### Genes differentially spliced in breast tissue are enriched in estrogen response pathway

The highest number of differentially spliced genes was observed in breast tissue. We reasoned that this gene set would have the highest statistical power to reveal enriched pathway associations. Analysis of this set using the compute overlaps tool of the Molecular Signatures Database (*48*) revealed estrogen response (`HALLMARK_ESTROGEN_RESPONSE_EARLY`) as the top enriched gene set, with a total of 17 associated genes (FDR = 1.8 × 10^−8^; table S10).

## Discussion

Our study has shown that males and females display widespread differences not only in gene expression but also in differential splicing. Gene expression is defined in our study based on the total read counts for all isoforms of a gene. Differential splicing is defined as a change in the level of one or more isoforms of a gene relative to the others. Our definition of significant alternative splicing as defined by equation (1) in the Methods identified a set of genes 70% of which were distinct from the genes found to be significantly differentially expressed.

Our results showed the highest density of significant DAS events to be on the X chromosome, which is similar to recent findings on sex-biased differential expression (*17*). We additionally showed that both differentially expressed and differentially spliced X chromosomal genes escape X-inactivation (Fig. 3). Interestingly, the number of genes showing sex-biased AS normalized by exon count was strikingly different also among the autosomes (Fig. 4).

We identified a small number of genes with sex-biased DAS across multiple tissues, but the vast majority of events (97.6%) were specific to one tissue. This contrasts with recent findings at roughly 18% of sex-biased expression was discovered in a single tissue (*17*). The lower proportion of DAS events detected in multiple tissues could be related to a lower statistical power of identifying DAS events compared to DGE events. Larger cohorts, as well as quantitative long-read transcriptome sequencing, could be helpful to further characterize tissue specificity of sex-biased splicing.

Our results suggest that a portion of sex-biased DAS events are driven by estrogen. Estrogen receptors *α* and *β* function as transcription factors that mediate the cellular effects of estrogen hormones, particularly the main circulating estrogen hormone 17*β*-estradiol (E2) by mechanisms including the activation of target genes (*49*). In addition, ER*α* and ER*β* can mediate estrogen-induced alternative splicing (*43, 50*). Our results do not allow conclusions about the molecular mechanism, however. ERs can directly interact with chromatin at specific DNA sequences known as estrogen response elements or indirectly regulate gene expression by means of a variety of intracellular signaling events and in some case may contribute to lasting epigenetic changes that in turn can influence gene regulation (*51, 52*). Functional genomic experiments are needed to characterize the mechanism of estrogen’s influence on sex-specific alternative splicing.

Our results suggest that future studies on the relationship of gene expression and disease should include an analysis of splicing by sex. The sex differences observed in our analysis were by design on largely healthy tissues of older individuals, and may not be representative of differences seen in various disease states. Interestingly, some of the sex-biased splicing events have also been observed in cancer, including those described above for *FGFR2* (*53*), *ESYT2* (*54*), and *SLK* (*55*), although future work will be required to determine if the sex-biased DAS observed here is related to sex differences in cancer. Our results provide an atlas of tissuespecific sex differences that can be used to interpret changes seen in disease settings.

Our study provides a comprehensive atlas of genes showing sex-biased AS across a range of tissues. AS is becoming more easily accessible thanks to long-read transcriptome sequencing (*56*), and we anticipate that more detailed investigations of sex differences in specific diseases will be valuable for advancing precision medicine for males and females.

## Methods

### GTEx samples

FASTQ files as well as transcript per million (TPM) and read counts of 56,202 genes together with the corresponding GTEx sample attributes and phenotypes were downloaded from the most current release, GTEx V8 (dbGaP Accession phs000424.v8.v2 (released 2019-08-26). Approval for use of the raw GTEx RNA-seq FASTQ files was granted by Database of Genotypes and Phenotypes (dbGaP). We restricted our analysis to tissues for which at least 50 samples were available for both males and females (Table 2).

**Table 2:** GTEx Samples. Samples from 351 males and 181 females were analyzed. Tissues were included in the analysis if at least 50 samples were available for each sex. A total of 39 tissues were included with 10444 male, 5087 female, and 15531 total samples.

| **Tissue** | **Male** | **Female** | **Total** |
| --- | --- | --- | --- |
| Adipose (sc) | 445 | 218 | 663 |
| Adipose (v) | 371 | 170 | 541 |
| Adrenal gland | 157 | 101 | 258 |
| Aorta | 279 | 153 | 432 |
| Coronary artery | 146 | 94 | 240 |
| Tibial artery | 454 | 209 | 663 |
| Caudate | 183 | 63 | 246 |
| Cerebellar hemisphere | 157 | 58 | 215 |
| Cerebellum | 174 | 67 | 241 |
| Cortex | 181 | 74 | 255 |
| Frontal cortex | 153 | 56 | 209 |
| Hippocampus | 143 | 54 | 197 |
| Hypothalamus | 147 | 55 | 202 |
| Nucleus accumbens | 182 | 64 | 246 |
| Putamen | 156 | 49 | 205 |
| Spinal cord | 102 | 57 | 159 |
| Breast | 291 | 168 | 459 |
| Fibroblasts | 330 | 174 | 504 |
| EBV-lymphocytes | 112 | 62 | 174 |
| Sigmoid colon | 240 | 133 | 373 |
| Transverse colon | 259 | 147 | 406 |
| Esophagus (gej) | 251 | 124 | 375 |
| Esophagus (m) | 363 | 192 | 555 |
| Esophagus (mu) | 338 | 177 | 515 |
| Atrial appendage | 293 | 136 | 429 |
| Left ventricle | 294 | 138 | 432 |
| Liver | 161 | 65 | 226 |
| Lung | 395 | 183 | 578 |
| Skeletal muscle | 543 | 260 | 803 |
| Tibial nerve | 419 | 200 | 619 |
| Pancreas | 207 | 121 | 328 |
| Pituitary | 204 | 79 | 283 |
| Skin (not exposed) | 411 | 193 | 604 |
| Skin (exposed) | 467 | 234 | 701 |
| Small intestine | 120 | 67 | 187 |
| Spleen | 154 | 87 | 241 |
| Stomach | 227 | 132 | 359 |
| Thyroid | 434 | 219 | 653 |
| Whole blood | 501 | 254 | 755 |
| **Total** | **10444** | **5087** | **15531** |

We used the Yet Another RNA Normalization software pipeline (YARN) to look for samples that are likely to be mis-annotated. We applied the function checkMisAnnotation using chromosome Y genes as control genes and removed the individual GTEX-11ILO from the dataset (*57*).

### Alignment of RNA-seq data

A matrix of counts for each of a variety of splicing types was generated by the rMATS (*58*) program (version 3.2.5) for each of the samples available from the GTEx archive. FASTQ files from the GTEx project were aligned to the Genome Reference Consortium Homo sapiens assembly version hg38 (GRCh38.p12) using hisat2 (*59*), and duplicates were removed using the Picard toolkit (http://broadinstitute.github.io/picard/). In order to create a matrix of counts with rows containing unique junction identifiers for each of the splicing types and columns containing the unique GTEx sample identifiers, some modifications were made to the standard process of running the rMATS program. For each file, rMATS identifies specific alternative splicing events capturing skipped exons (SE), retention introns (RI), alternative 3’ and 5’ splice sites (A3SS and A5SS), and mutually exclusive exons (MXE). For each of these 5 different splicing types, rMATS creates two files, one containing the counts of reads that span the splicing junctions only, and a second file that additionally contains the counts of reads that are on target.

Data from individual samples were merged into a single matrix for each of the sample types and for each of the AS types.

### Differential gene expression in male vs. female tissues

For analysis of differential gene expression and alternative splicing, genes on the Y chromosome were excluded from the analysis. In addition, for each tissue we kept only events for which the number of male and female samples with at least 1 cpm (count per million) for the gene of interest exceeds a threshold *X*, where *X* = 0.25·min(*M, F*), where *M* is the number of male samples for a particular tissue, and *F* is the number of female samples. For each tissue, genes differentially expressed between male and female were individually determined using the voom function from the R package limma (*60, 61*).

### Characterization of alternative splicing events (ASEs)

We investigated individual AS events rather than transcript (isoform) abundance, because despite improvements in algorithms, accurate quantification of the expression of individual transcripts is challenging with short-read RNA-seq technology, especially for short or low-abundance transcripts and genes with complex structures (*62–65*).

We used rMATS to identify and count reads that correspond to each of the 5 types of ASEs: (1) skipped exon - the skipping of a single exon in an isoform of the transcript. (2) mutually exclusive exons - two consecutive exons out of which only one is present in each isoform of the transcript. (3) retained intron - the retention of an intron in an isoform of the transcript. (4) alternative 5’ splice site - a different exon at a 5’ position in an isoform of the transcript. (5) alternative 3’ splice site - similar to the previous category, but at a 3’ position (See Supplemental Fig. S2). rMATs identifies these events from a GTF file of known transcripts (for the experiments described here, release 25 from GENCODE annotation for genes for GRCh38.p7 was used). rMATS then counts the number of reads that agree with each of the two alternatives that the event describes. For example, for a skipped exon rMATS will count the reads that fall within splice junctions that connect the skipped exon to its neighboring exons, and the reads that fall within a splice junction that connects the neighboring exons to each other. A matrix of event counts was generated for all samples according to tissue types; one matrix was generated for each of the 5 categories of ASE and was used for the downstream analysis.

### Statistical approach to differential splicing between males & females

In order to be able to fit a linear model to the data we used voom (*60*) to transform the counts into continuous data, appropriate for linear modeling. For each ASE, we combine skip and inclusion event counts as individual samples in a multifactorial linear model, where skip and inclusion counts from the same individual/sample are treated as replicate arrays in order to account for correlation. Limma uses generalized least squares to fit the model, which does not assume that the errors of different samples are independent. Specifically, we pass to the functions voom and lmFit the block and correlation arguments, where block encodes to the replicate structure and correlation is obtained with the limma function duplicateCorrelation(). The multifactorial model has 3 predictors: sex (male or female), event (skip or inclusion) and a sex:event interaction term:

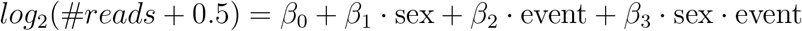

Events that have a significant `sex · event` interaction term (FDR≤0.05), and in addition a fold change of at least 1.5 for that term, are considered differentially spliced. The sex predictor accounts for the case where males or females have a higher level of both isoforms but the proportions in both sexes are the same. The event predictor accounts for the case where one event has more reads mapped to it, but not as a result of alternative splicing that is differential between the sexes. For example, if due to the fragmentation process of RNA-Seq more reads are mapped to the inclusion event, there will be a bias in both sexes towards this event. For normalization, we used the edgeR function `calcNormFactors` applying TMM normalization (*66, 67*). An explanatory cartoon is shown in Supplemental Fig. S3.

We kept only events with at least 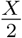 male samples had a cpm ≥ 1 and analogously for the female samples in each tissue, where *X* equals the size of the smallest study group (male or female). This was done to prevent genes displaying nearly only inclusion or exclusion isoforms from falsely being called differentially spliced (see Supplemental Fig. S4 for an example).

### Calculation of the Sex-Biased Splicing Index

The normalized sex-biased splicing index is defined as the number of statistically significant splicing events per 1000 exons in the chromosome. The normalized splicing index equals the number of exon skipping events divided by the number of exons on the chromosome and multiplied by 1000.

### Identification of events included in a nonsense-mediated decay isoform

For each skipped exon event, we find all isoforms that contain this exon in the annotation file gencode.v30.annotation.gtf. For each isoform, we compare the amino acid length of the inclusion isoform to the sum of lengths of the skip isoform and the skipped exon. If the former is smaller, then the inclusion isoform is classified as an NMD isoform containing a premature truncation codon as a consequence of including the exon. To obtain the lengths of each isoform, we used the program gffread from the GffCompare package (*68*).

### Isoform-level analysis of breast cell lines exposed to estradiol

Paired-end fastq files were downloaded from the SRA, quality-controlled using fastp (*69*), and aligned using STAR (*70*) via RSEM (*45*), following by isoform quantification and differential splicing prediction by HBA-DEALS (*46*). For each cell line, isoforms that are expressed in all samples were considered expressed, and genes that had at least 2 expressed isoforms were analyzed. The most significant event/isoform of a gene was used to determine if a gene is DAST. Hypergeometric p-values were computed for genes that were present (analyzed) in both datasets, and combined using Fisher’s method. For HBA-DEALS, to determine whether the covariates alpha and beta are in the same layer or in separate layers we computed the Bayes factor for a sample of genes and followed the majority vote. To compute the p-values we used the region of practical equivalence (*71*), with magnitude 0.1 for expression and the same value scaled by the denominator of the Aitchison perturbation for splicing. For multiple testing correction, we created a model for all the gene together, set a mixture prior for expression (Gaussian) and splicing (Dirichlet) such that one component is weakly informative (as in the non-crrected model), and one is concentrated within the region of practical equivalence, and found the mixture weights that maximize the joint posterior using the rstan function ‘optimizing’. We then set these values as fixed mixture weights and use the same mixture priors in finding the posterior of each gene separately (*72*). Isoforms were classified as DAST using a corrected threshold of 0.1. All of these functionalities are implemented in the HBA-DEALS R package, which is freely available at https://www.github.com/TheJacksonLaboratory/HBA-DEALS

### Overlap of introns of alternatively-spliced events and estrogen receptor ChIP-Seq peaks

The bed file with ESR1 peaks corresponding to the dataset of Ross-Innes et al. (*47*) was downloaded from the Cistrome database (*73*). Overlap between introns and peaks was calculated using bedtools ().

### Gene Ontology analysis

For each of the tissues, Gene Ontology (GO) term enrichment analysis was performed. The study set was defined as the set of all significantly differentially spliced genes in the tissue. For comparison, analogous experiments were performed with differentially expressed genes. The population set was defined to be all genes included in the analysis of expression or splicing. Analysis was performed using the Term-for-Term approach with an updated version of the Ontologizer (*74*) as implemented in Phenol (https://github.com/monarch-initiative/phenol). p-values were adjusted for multiple testing using the Bonferroni method.

### Code availability and reproducibility

A pipeline to retrieve RNA sequencing data from SRA, perform alignment with hisat2, and characterize splicing events with rMATS 3.2.5 was developed using the Nextflow workflow framework (*75*). The workflow is publicly available in GitHub at https://github.com/lifebit-ai/rmats-nf. To enable the execution of this large scale analysis of a total of 7906 SRA samples, we executed the Nextflow analyses as batches of up to 500 samples. The analyses were performed in the Lifebit CloudOS platform and we have made available the analysis page shareable links in Supplementary Table S11.

A companion Nextflow workflow was also developed to facilitate the collation of the rMATS derived matrices from the individual batches into 50 final tables and can be accessed at https://github.com/lifebit-ai/merge-rmats-nf. These Nextflow analyses were also run in the Lifebit CloudOS platform and the shareable analysis links are provided in (that Supplementary table).

We developed custom scripts, in the Jupyter notebook format, to implement the analyses for the sections mentioned above in Methods, “Differential gene expression in male vs. female tissues”, “Characterization of alternative splicing events (ASEs)”, “Statistical approach to differential splicing between males & females”, “Calculation of the Sex-Biased Splicing Index”, and “Identification of events included in a nonsense-mediated decay isoform”. All the Jupyter notebooks are publicly accessible at https://github.com/TheJacksonLaboratory/sbas/tree/master/jupyter.

To enable the reproduction of the results from the notebooks, we have made available the input data via a Zenodo record, that is publicly accessible at https://zenodo.org/record/4179559. To facilitate the reproduction of the results, we have created a helper bash script, that programmatically executes the jupyter notebooks, leveraging the papermill library (https://github.com/nteract/papermill). The helper script can be found at https://github.com/TheJacksonLaboratory/sbas/blob/master/reproduce.sh.

## Acknowledgments

Support for this work was provided by the National Institutes of Health, through the Data Commons program (award 1OT3OD025646-01 [Helium]) and by the Donald A. Roux family fund. O.A. was supported by The Jackson Laboratory Start Up Funds.

The Genotype-Tissue Expression (GTEx) Project was supported by the Common Fund of the Office of the Director of the National Institutes of Health (commonfund.nih.gov/GTEx). Additional funds were provided by the NCI, NHGRI, NHLBI, NIDA, NIMH, and NINDS. Donors were enrolled at Biospecimen Source Sites funded by NCI Leidos Biomedical Research, Inc. subcontracts to the National Disease Research Interchange (10XS170), Roswell Park Cancer Institute (10XS171), and Science Care, Inc. (X10S172). The Laboratory, Data Analysis, and Coordinating Center (LDACC) was funded through a contract (HHSN268201000029C) to the The Broad Institute, Inc. Biorepository operations were funded through a Leidos Biomedical Research, Inc. subcontract to Van Andel Research Institute (10ST1035). Additional data repository and project management were provided by Leidos Biomedical Research, Inc.(HHSN261200800001E). The Brain Bank was supported supplements to University of Miami grant DA006227. Statistical Methods development grants were made to the University of Geneva (MH090941 & MH101814), the University of Chicago (MH090951,MH090937, MH101825, & MH101820), the University of North Carolina - Chapel Hill (MH090936), North Carolina State University (MH101819),Harvard University (MH090948), Stanford University (MH101782), Washington University (MH101810), and to the University of Pennsylvania (MH101822). The datasets used for the analyses described in this manuscript were obtained from dbGaP at http://www.ncbi.nlm.nih.gov/gap through dbGaP accession number phs000424.v8.v2.

## Author Contributions

G.K. and P.N.R. conceived of and designed the study. D.F.T.V., A.D.M., C.C., P.P.B., A.K.K., D.D., X.A.Z., J.G., R.S., P.H., D.S., J.Y. developed software used in the analysis, G.K. and P.N.R. wrote the manuscript with contributions from J.A.M., M.A.H., G.K., T.O., O.A., and J.B.; P.N.R. supervised the study.

