## Supplementary material for "The impact of biological sex on alternative splicing": Suppl. Figures and Tables

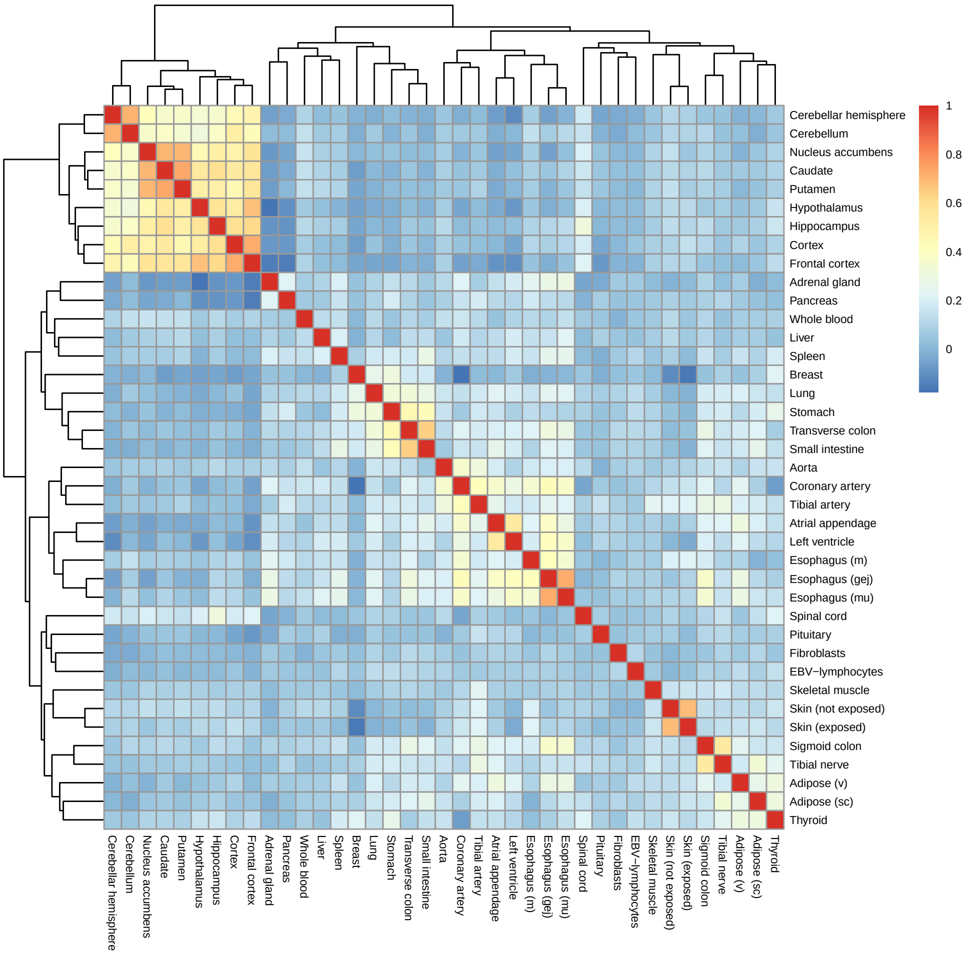


**Figure S1: Heatplot representing similarity in the fold-changes between male and female samples**. The heatplot represents a hierarchical clustering of the different tissues, where tissues are clustered by similarity of the mean fold changes of gene expression between male and female samples.

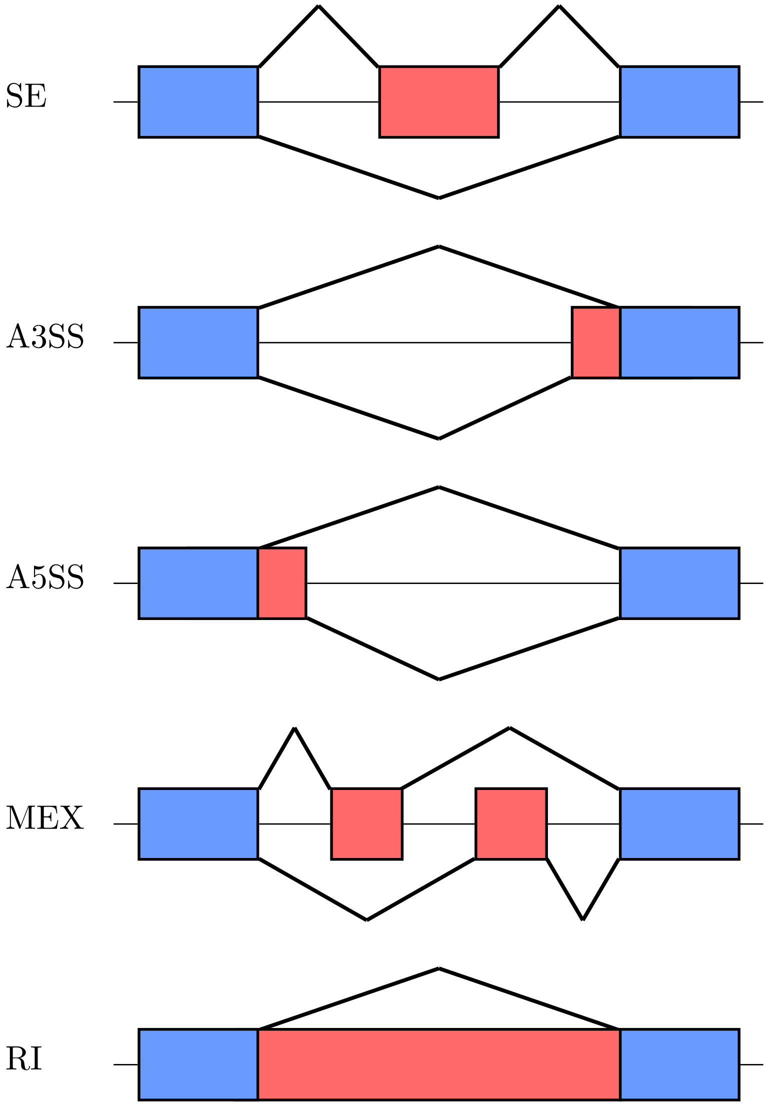


**Figure S2: The five categories of AS events investigated in this work** SE: skipped exon/exon inclusion; A3SS/A5SS: alternative 3'/5' splice site; MEX: mutually exclusive exons; RI: retained intron.

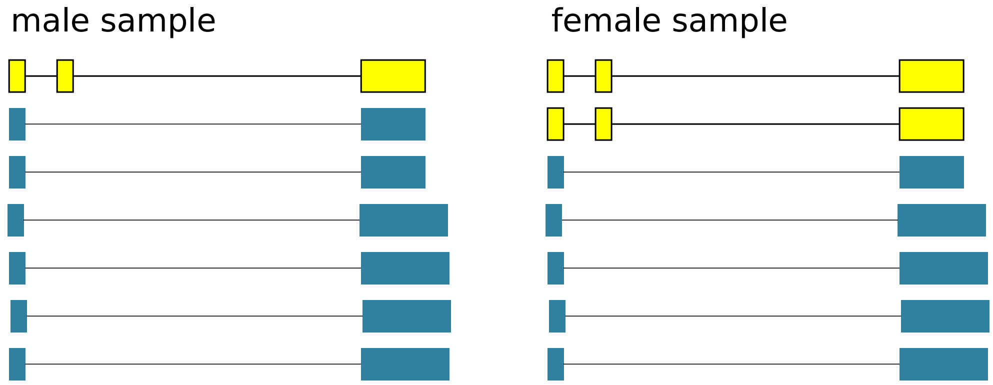


**Figure S3: Linear model for sex-biased alternative splicing.** In this example, we are focused on a gene for which 7 transcripts are sequenced in the male sample and 7 in the female sample. Concentrating on the middle exon of the transcript on the top line, the male inclusion count is 1, the male exclusion count is 7, and the female inclusion count is 2 and the female exclusion count is 6. The regression is a linear model in which the number of reads is predicted separately for inclusion and exclusion reads as a function of a baseline ($\beta_0$), sex ($\beta_1$), with female encoded as 0 and male as 1, the alternative splicing event ($\beta_2$) encoded as 0 for skipping and 1 for inclusion (and analogously for other types of AS), and finally an interaction term reflecting the influence of sex on the event counts ($\beta_3$). We called AS events significantly sex-biased if the interaction term was significant following multiple testing correction.

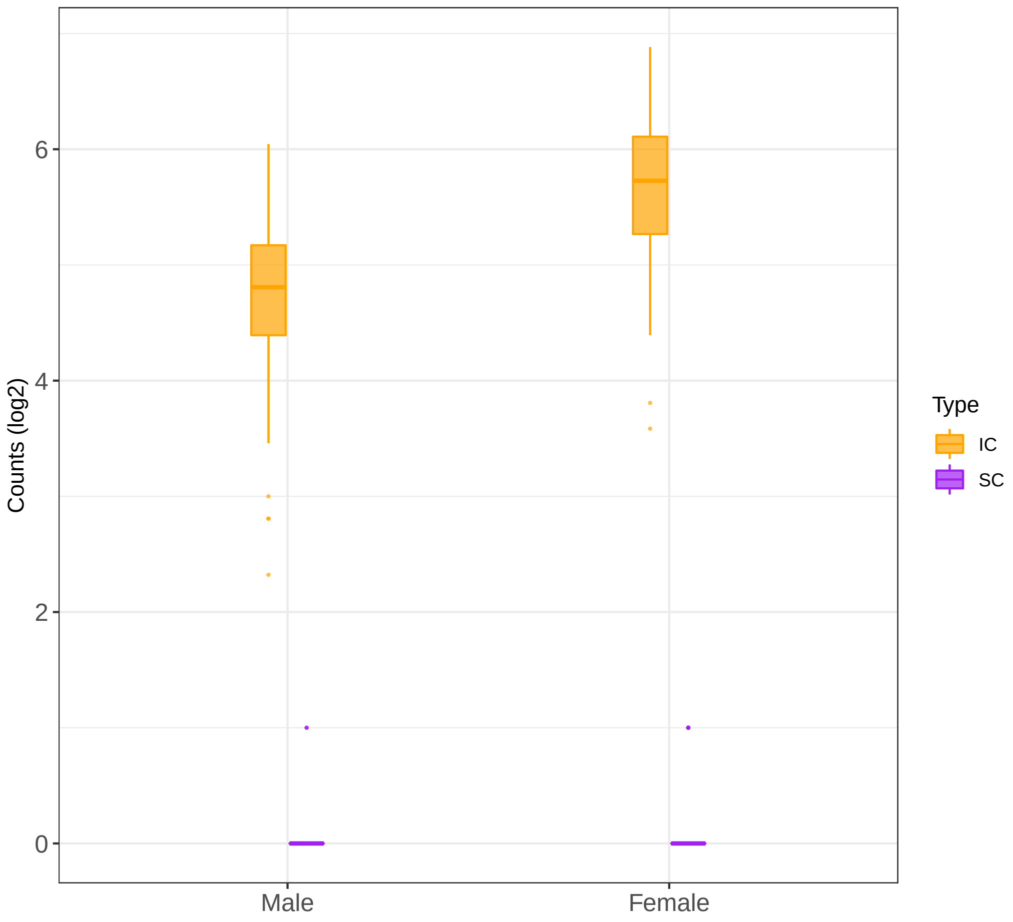


**Figure S4: Usage of just one of two potential isoforms.** The box plot shows *KDM5C* counts for an event with only two isoforms (one inclusion and one skipped exon). The skipped isoform is not expressed either in females and males. Even though there is a significant difference in the inclusion counts, we do not interpret this as differential splicing, because only one form of the isoforms is expressed. This is the motivation for disregarding events for which less than $\frac{X}{2}$ male or female samples had a cpm $\geq 1$, where $X$ equals the size of the smallest study group (male or female). See the Methods of the main manuscript for details.

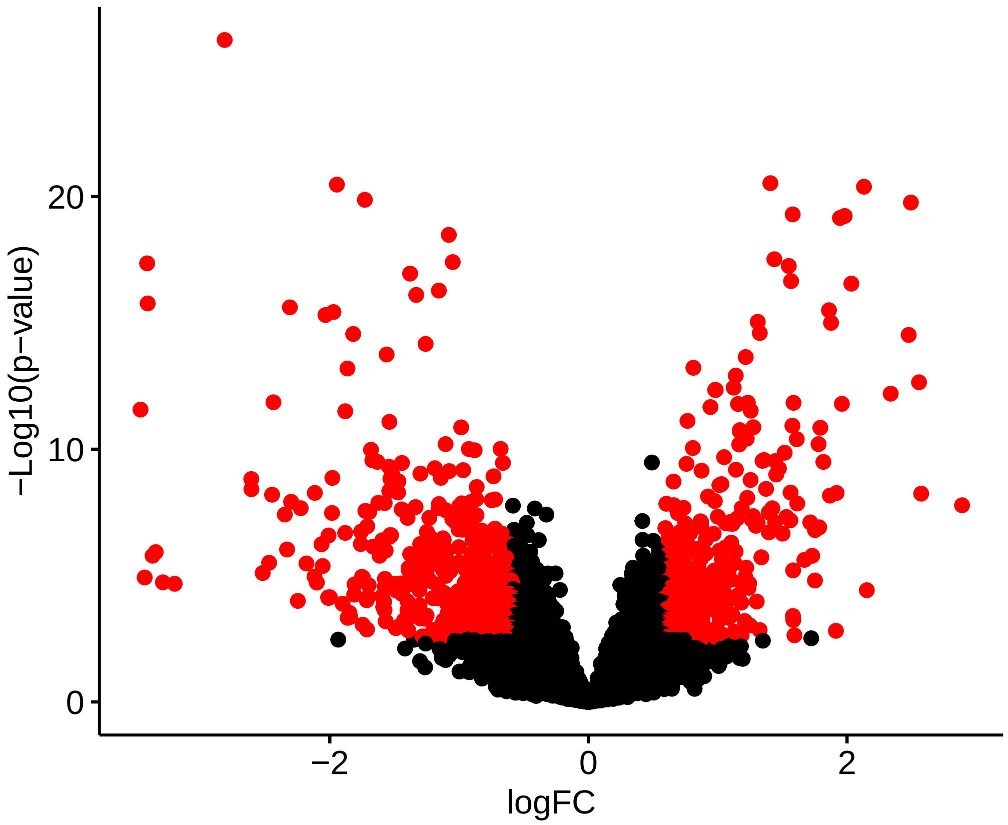


**Figure S5: Volcano plot of differentially spliced events in Breast -Mammary Tissue.** The log-fold-changes correspond to the change in inclusion read counts between males and females after accounting for differences that are due to differential expression or preferential mapping to one event type. The interaction term sex*isoform in the linear model captures changes in one isoform between males and females that are not explained by difference in expression (which are accounted for by the sex term) or preferential mapping of reads to one isoform type (which are accounted for by the isoform term). For example, if the number of skip counts are doubled in males compared to females, and this change is not due to random variation, but there isn't any significant difference in inclusion counts, then the gene is not differentially expressed, and the difference is not due to preferential read mapping. In this case we will classify the event as differential splicing event.

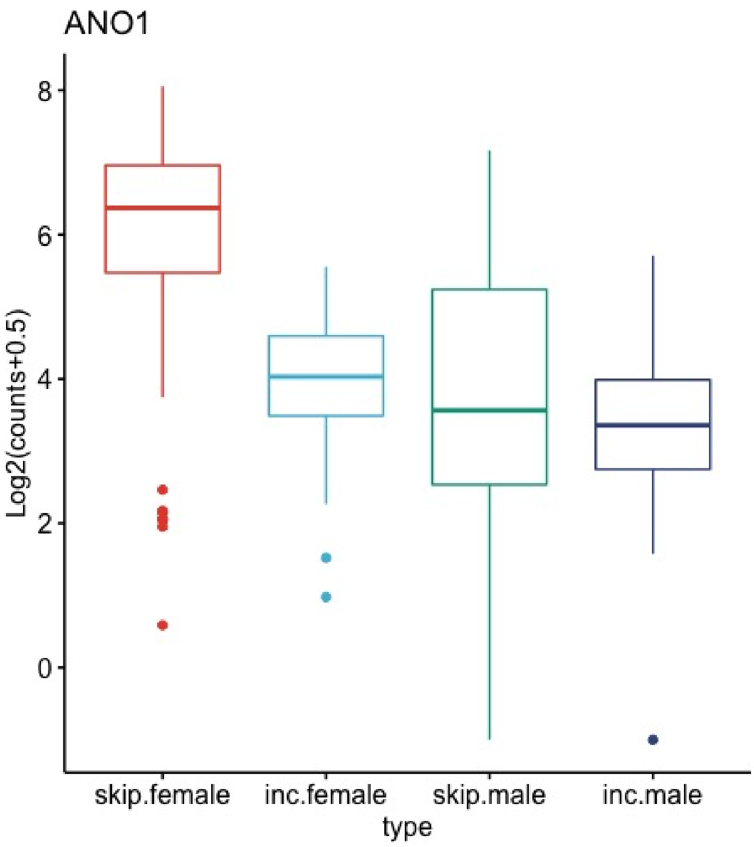


**Figure S6: ANO1.** Skipped exon event in *ANO1* in breast tissue (adjusted $p = 1.34 \times 10^{-8}$) The affected exon was ENSE00001470960 (chr11:70,155,911-70,155,988), length=78 nt (26 amino acids). ANO1 is a calcium-activated chloride channel. Splice variants of ANO1 have been shown to influence the biophysical properties of conductance [1]. The exon affected by skipping is exon 15 in Transcript ANO1-202 (ENST00000355303.9); it is located in the first intracellular loop within about 20 amino acids from a 9 amino acid segment that has been shown to that was shown to be crucial for both Ca(2+) and voltage sensing (cf. ref. [2] and Figure S1 therein). ANO1 modules EGFR- and CAMK-dependent pathways including AKT and MAPK [3]. Figure S8 shows the location of the skipped exon in the protein. shown in Figure S7. ANO1 overexpression in breast cancer is observed due to 11q13 amplification; anomalies of the distribution of ANO1 isoforms have been observed in breast cancer [4, 5].

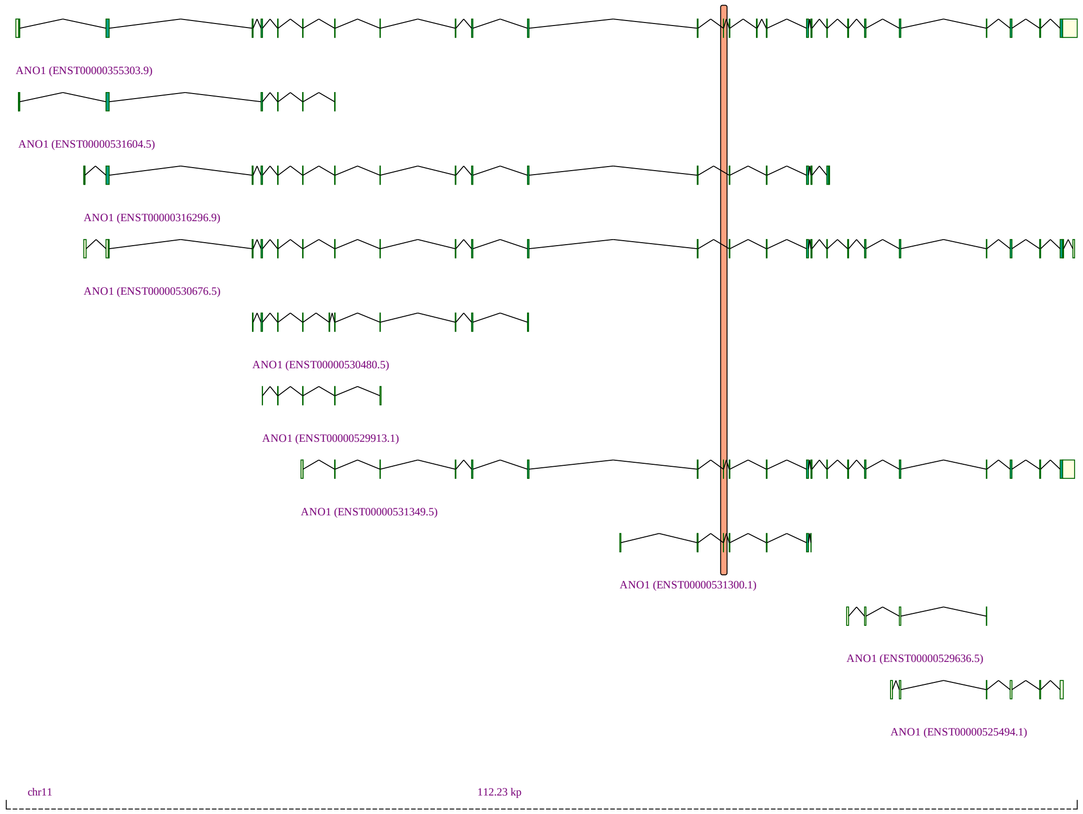


**Figure S7: ANO1.**

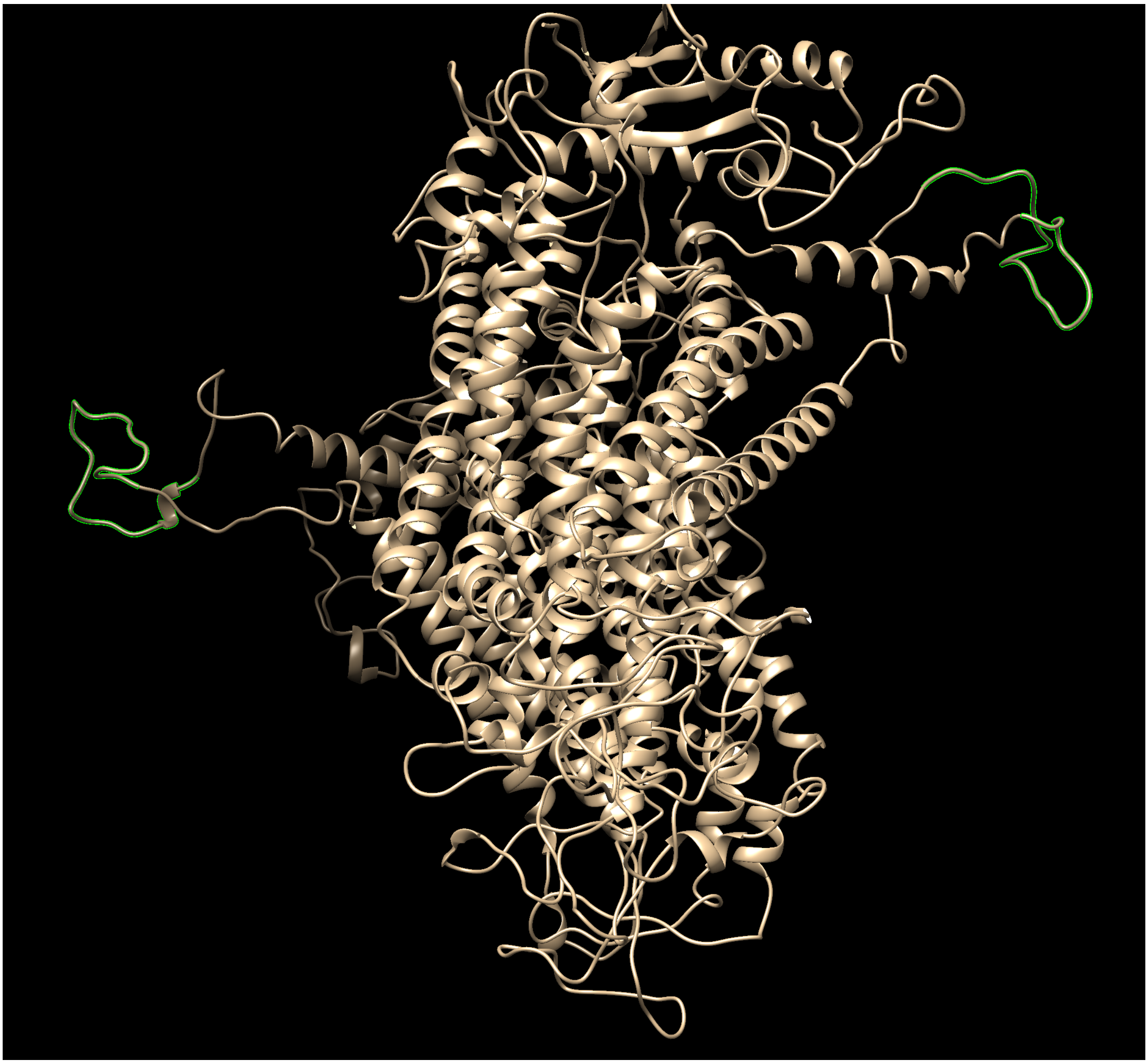

**Figure S8: ANO1.** Homology model of ANO1 protein structure (a dimer), with the region that is being removed by the skipping event of Fig. S7 highlighted in bright green. Graphic generated with the UCSF Chimera tool[6], structure generated using the Swiss-Model homology modeling server[7] .

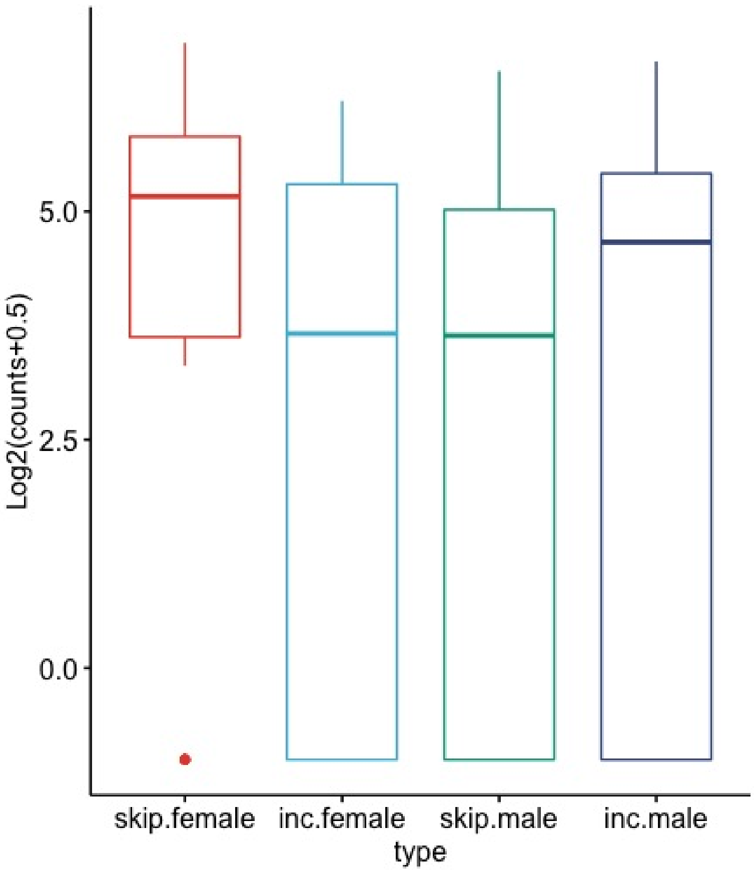


**Figure S9: AP3SL** Mutually exclusive exon event affecting AP3SL in hippocampus (adjusted $p = 0.013$). Adaptor related protein complex 3 subunit sigma 2 (AP3S2) is part of a protein complex that functions as a non-clathrin-associated adaptor complex in vivo [8]. The two exons are: (i) ENSE00001674538 (chr15:89,877,424-89,877,306). All transcripts that include this exon are annotated as nonsense-mediated decay (ENST00000423566.6, ENST00000423566.6, ENST00000558999.5, ENST00000558999.5, ENST00000560251.1, ENST00000560251.1); and (ii) ENSE00002551730.1 (chr15:89,878,300-89,878,265), which is included in one protein-coding (ENST00000558011.5) and one NMD (ENST00000558999.5) transcript. In Figure S10, ENSE00001674538 is highlighted in blue, and ENSE00002551730 is highlighted in orange. ENSE00001674538 displays more exon skipping in females. Since all isoforms that include this exon are NMD isoforms, but some isoforms that include the other exon are protein coding, this could indicate a partial shift away from NMD isoforms in females.

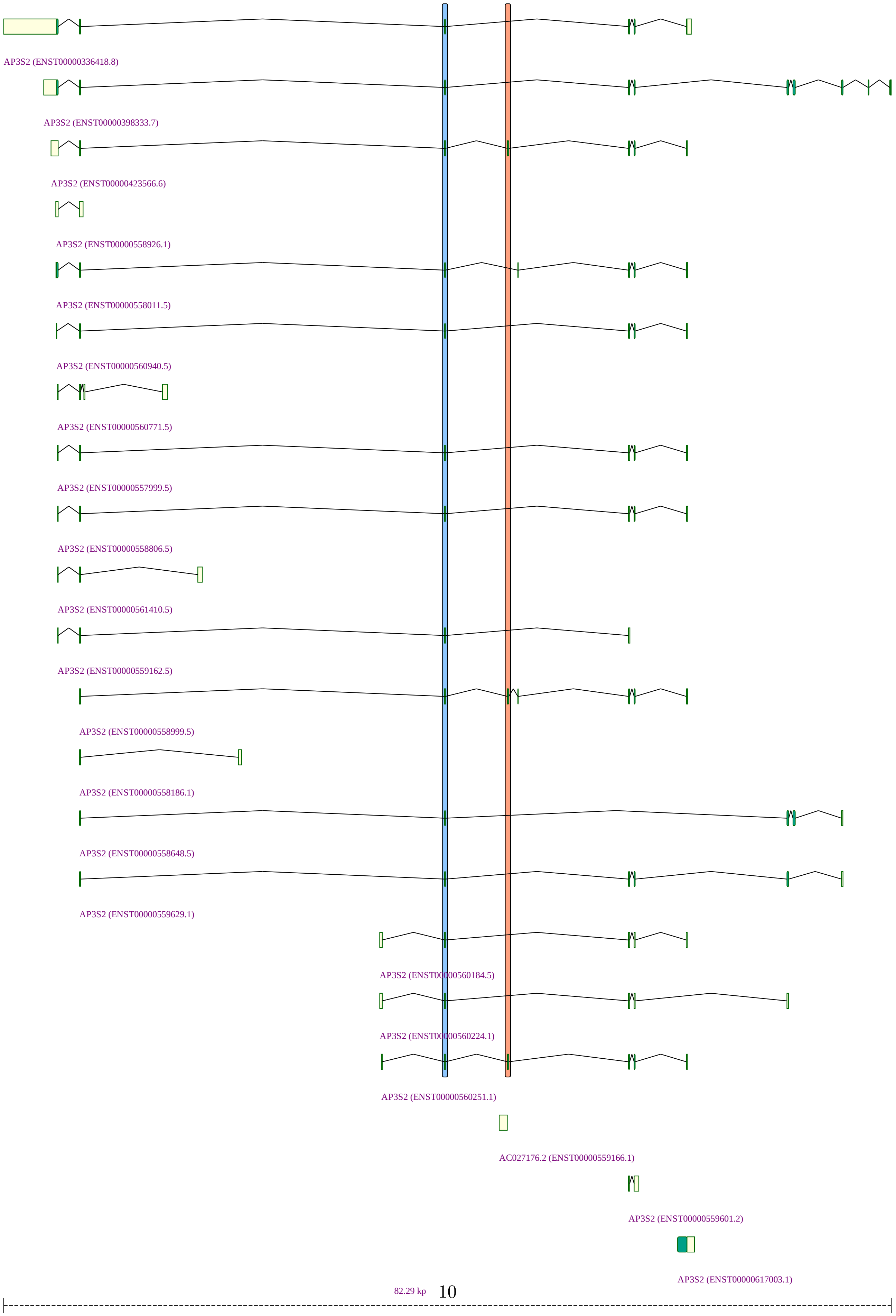




**Figure S10: AP3SL**

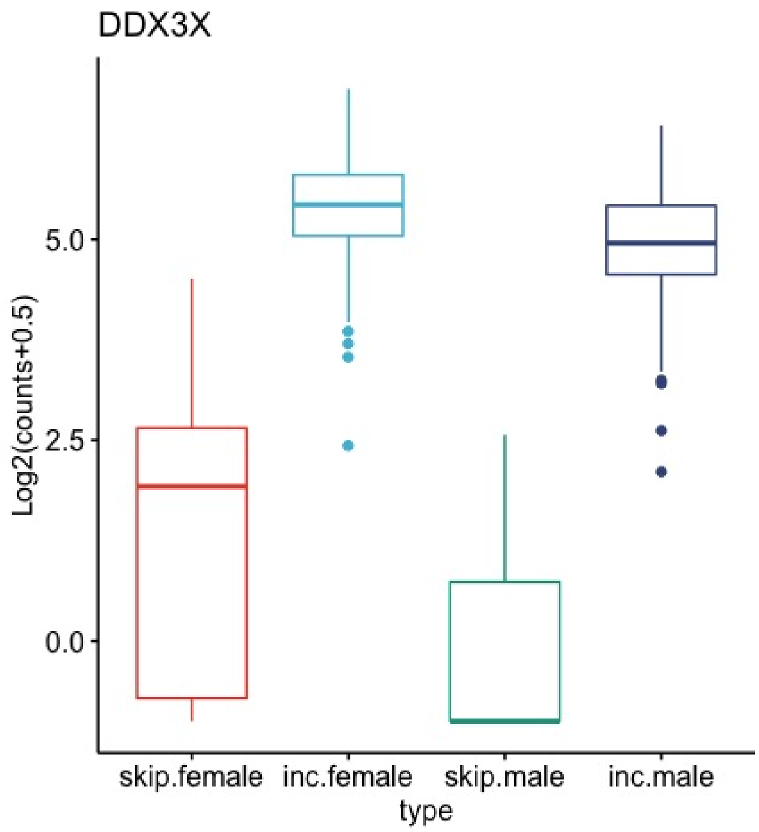


**Figure S11: DDX3X** *DDX3X* skipped exon event in skeletal muscle (adjusted $p = 6.23 \times 10^{-15}$). DDX3X is an RNA helicase with a DEAD/H (Asp-Glu-Ala-Asp/His) motif. DDX3X is involved in a variety of cellular biogenesis processes, including cell-cycle regulation, cellular differentiation, cell survival, and apoptosis [9]. The event affects exon ENSE00003458274.1 (chrX:41339038-41339083). Multiple isoforms include this exon, in some cases with the indicated boundaries differing by a few nucleotides). Figure S12 shows the location of the exon. Information about the function of the skipped isoform was not available in the literature at the time of this writing. The exon encodes part of the helicase ATP-binding domain [10], and speculatively, the exon skipping event could influence the function of that domain.

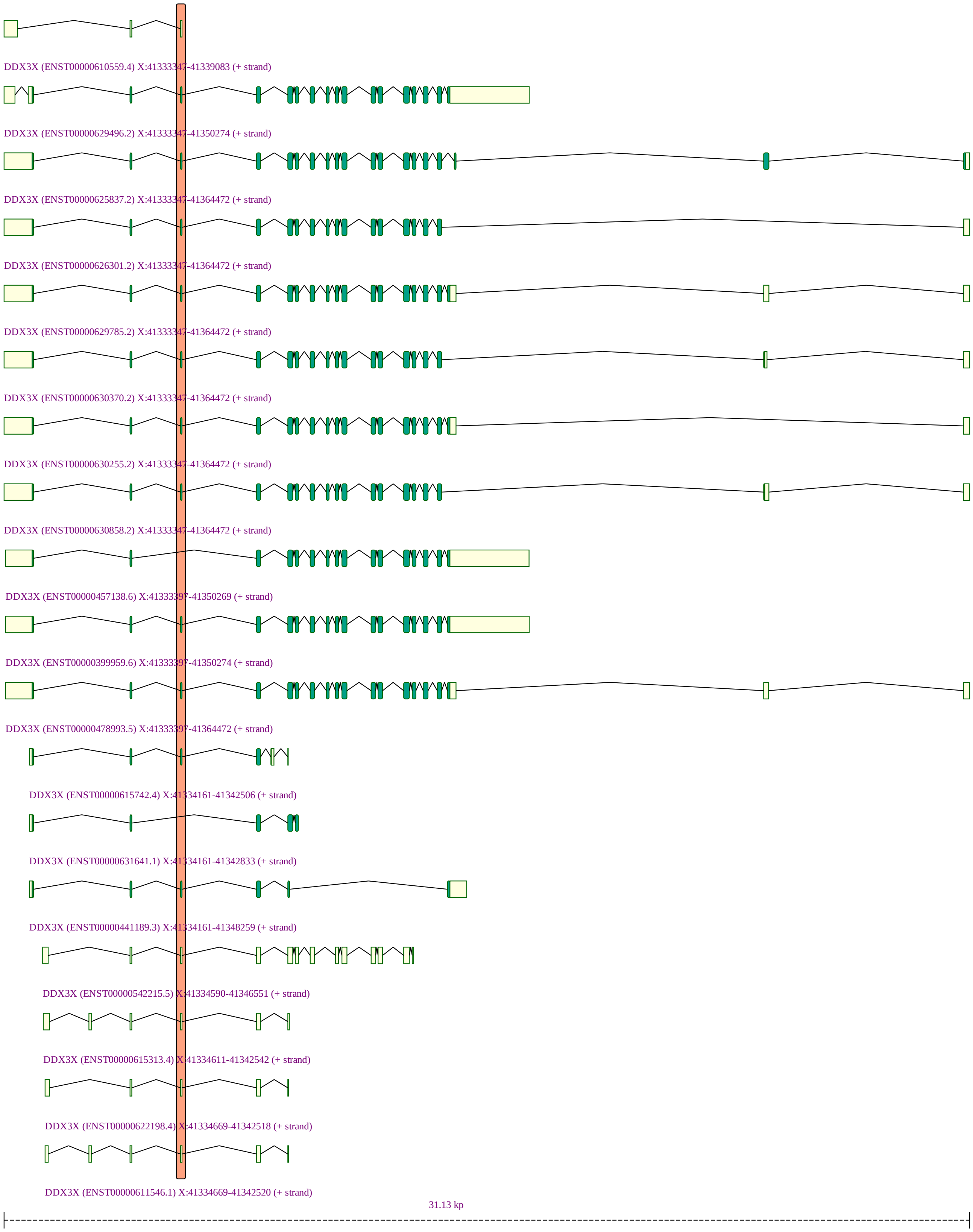


**Figure S12: DDX3X**

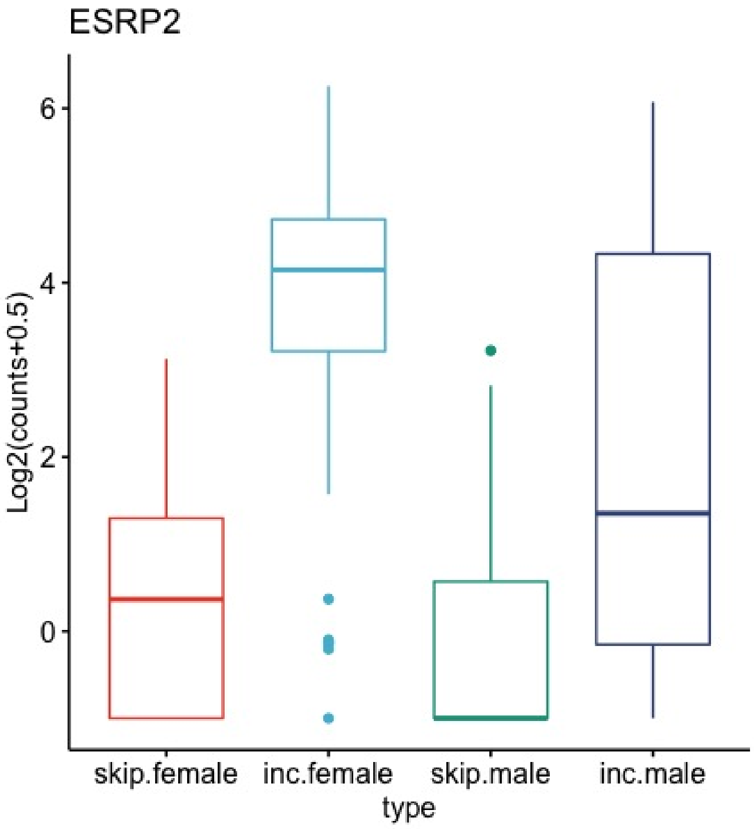


**Figure S13: ESRP2** *ESRP2* skipped exon event in breast tissue (adjusted $p = 6.97 \times 10^{-4}$). Epithelial Splicing Regulatory Protein 2 (ESRP2) is an epithelial cell-type-specific splicing regulator [11]. A skipped exon event was identified in breast tissue, with females showing 3 times more inclusion than skipping, a greater difference than males. The affected exon is ENSE00000691931.1 (chr16:68232370-68232503). ESRP2 may be able to regulate FGFR2 splicing [11]. Figure S14 shows the location of the affected exon.

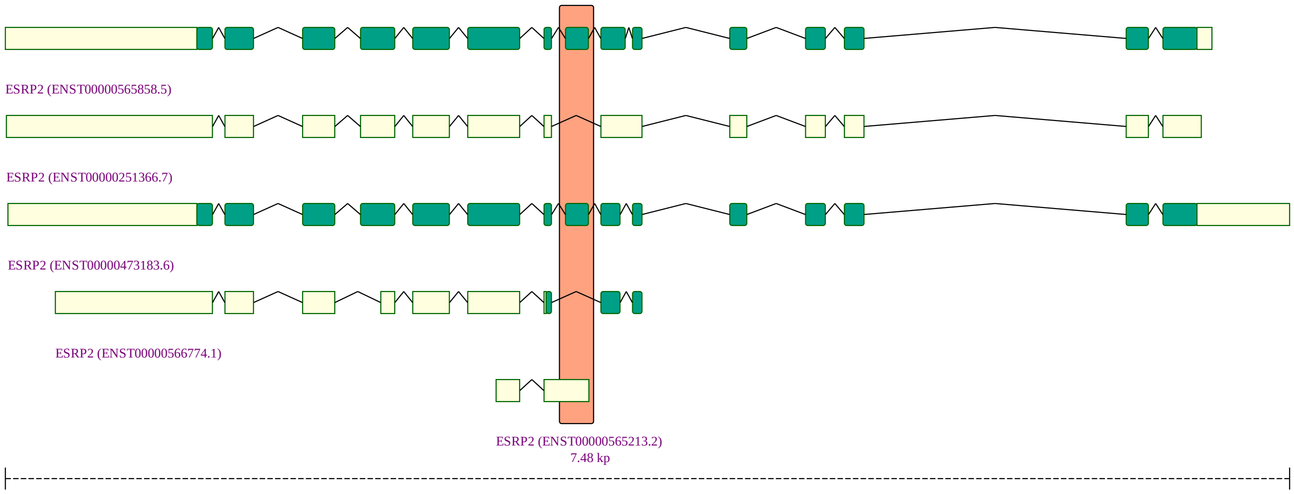


**Figure S14: ESRP2**

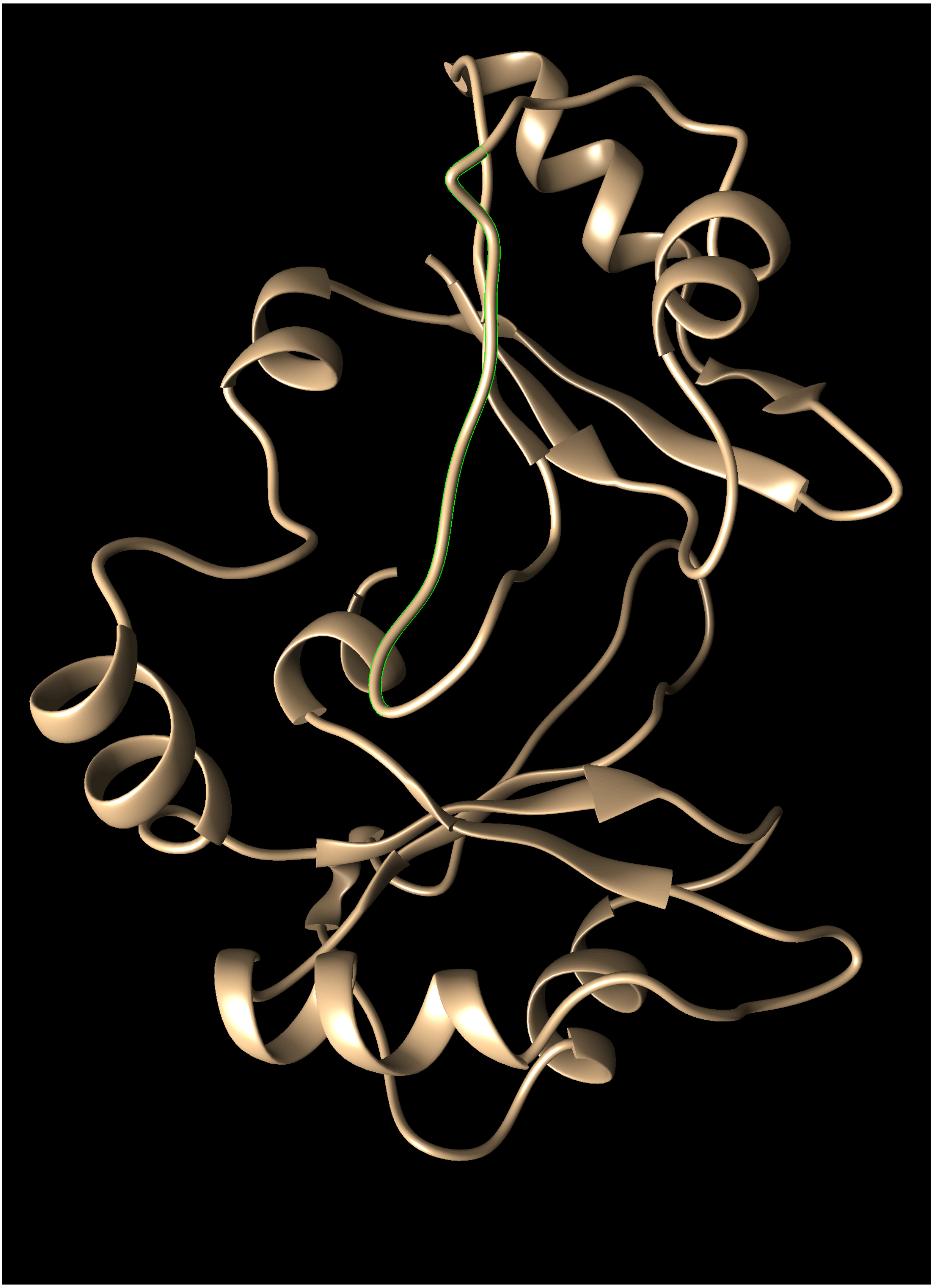

**Figure S15: ESRP2** Tertiary Structure of an ESRP2, with the region corresponding to the skipped-exon event highlighted in bright green. The structure was obtained using homology modeling[7]. Graphic generated with the UCSF Chimera tool[6]

.

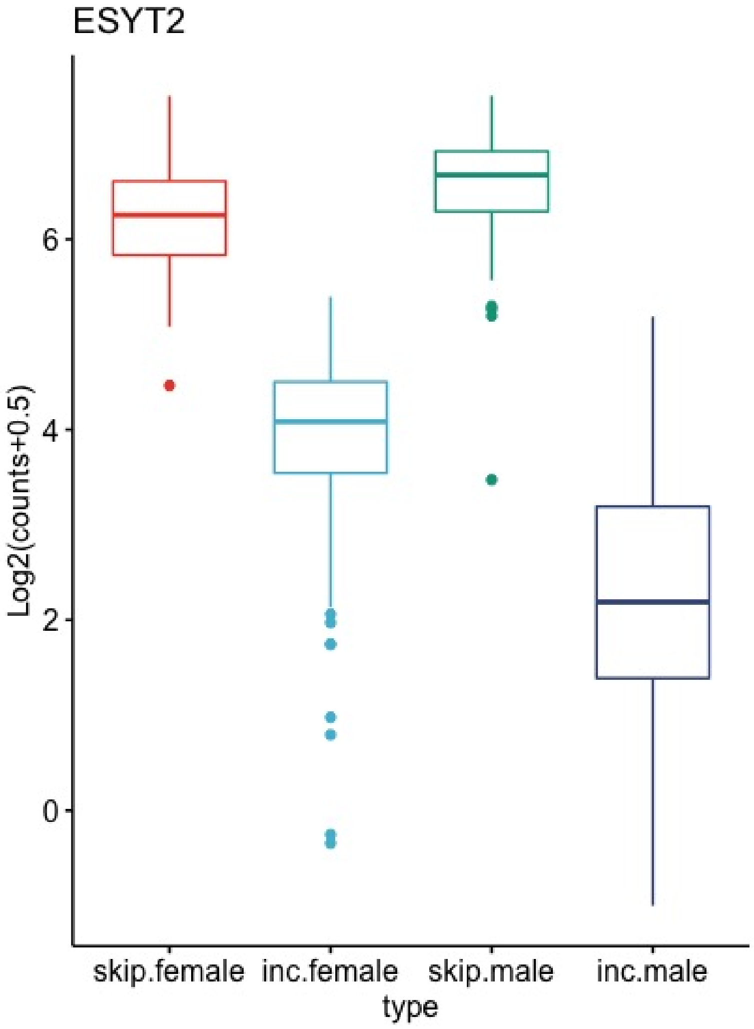


**Figure S16: ESYT2** Skipped exon event in *ESYT2* in breast tissue (adjusted $p = 2.22 \times 10^{-17}$). ESYT2 (Extended synaptotagmin-2) tethers the endoplasmic reticulum to the cell membrane and promotes the formation of appositions between the endoplasmic reticulum and the cell membrane, thereby acting as an ER-plasma membrane tether. Females show more inclusion of exon ENSE00001704685.1 (chr7:158752780-158752843). The inclusion of this 63 bp exon results in 21 additional amino acids in the C2B domain. Short and long isoforms of ESYT2 were shown to have differential ability to interact with stromal interaction molecule 1 (STIM1) [12].Figure S17 shows the location of the affected exon. Ca2+ release-activated Ca2+ (CRAC) channels mediate a sustained increase in cytoplasmic Ca2+ concentration essential for T cell activation. T cell receptor stimulation induces the depletion of the endoplasmic reticulum (ER) Ca2+ stores that activates CRAC channels via a process called store-operated Ca2+ entry (SOCE). STIM1, an ER-resident regulatory subunit that senses depletion of the ER Ca2+ stores, is an essential component of CRAC channels. Upon store depletion, STIM1 multimerizes and translocates from the ER to the ER-PM junctions where it promotes the opening of ORAI1, the pore subunit of CRAC channels. Among the E-Syt2 isoforms, only E-Syt2S supported STIM1 translocation to the junctions, and is uniquely abundant in T cells [12]. Lung tumors tend to express higher levels of the long splice variant (ESYT2-L), whereas the short variant (ESYT2-S) is the predominant form in non-malignant lung tissue. Knockdown of ESYT2-L in A549 cells or H2009 cells did not cause major changes in cytoskeleton organization, but knockdown of ESYT2-S caused a number of cytoskeletal abnormalities including a loss of $\alpha$-tubulin polarity [13]. These results suggest important functional differences between the two isoforms.

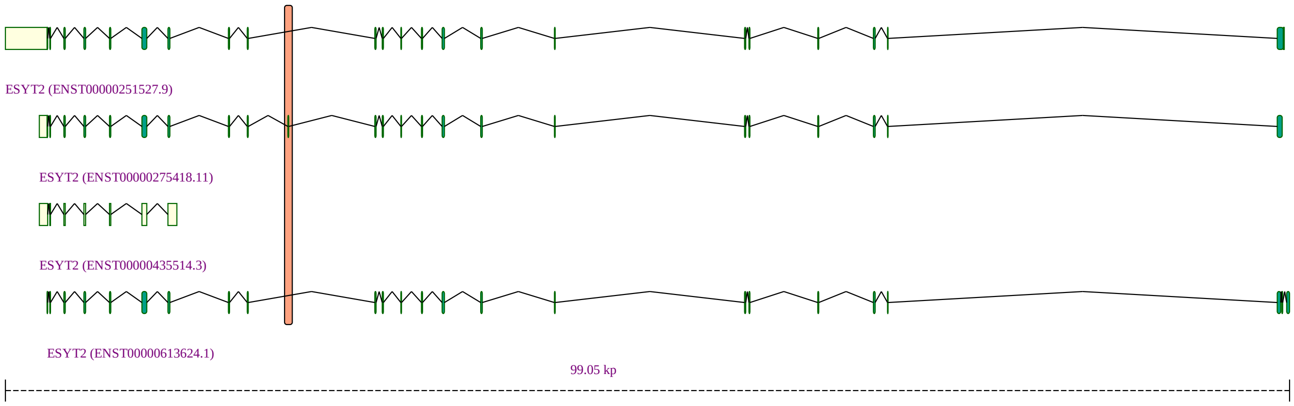


**Figure S17: ESYT2**

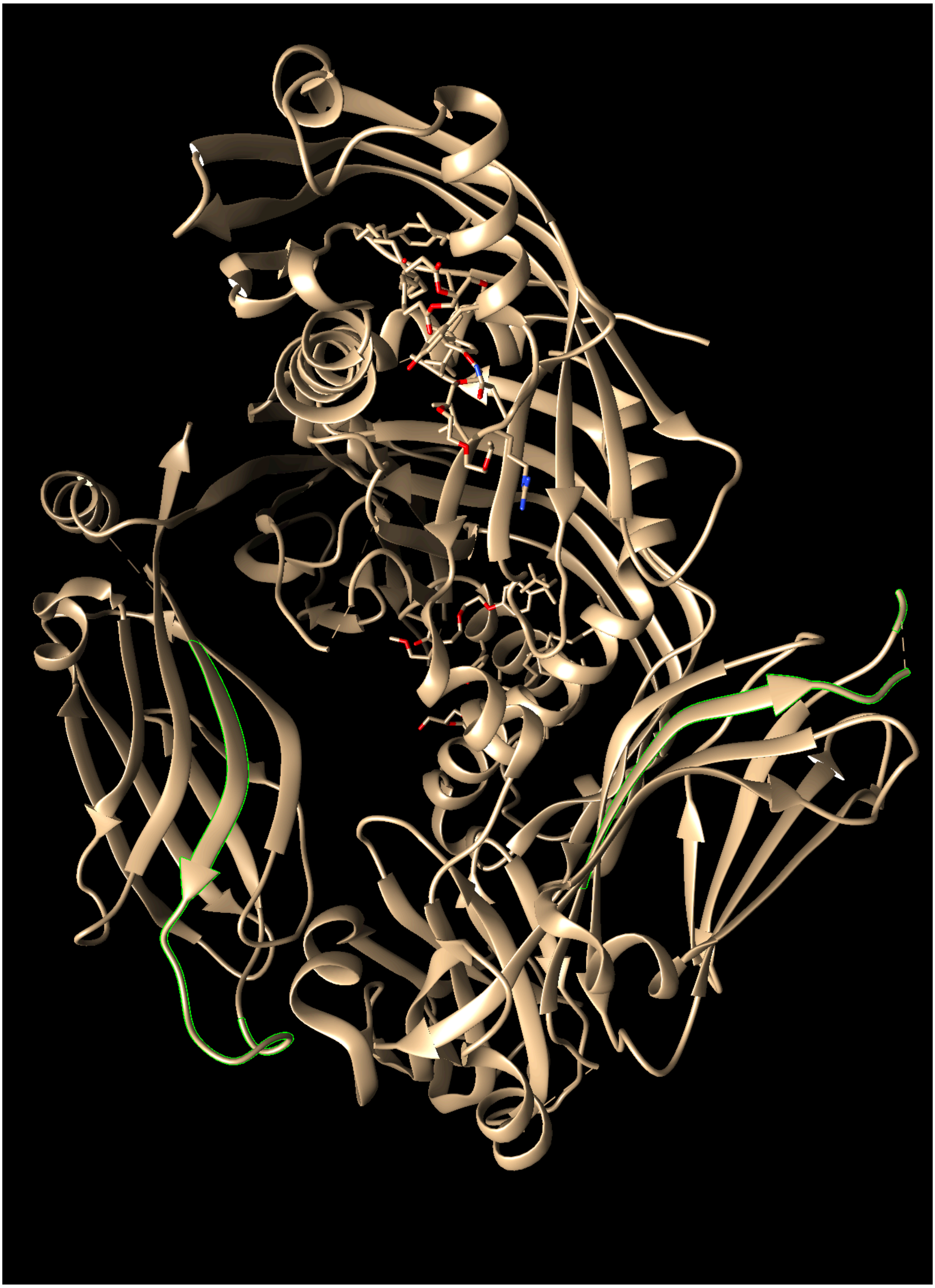

**Figure S18: ESYT2** Tertiary Structure of an ESYT2 dimer (skip isoform), with the region proximal to the skipping event highlighted in bright green (PDB ID 4P42). Graphic generated with the UCSF Chimera tool[6]
.

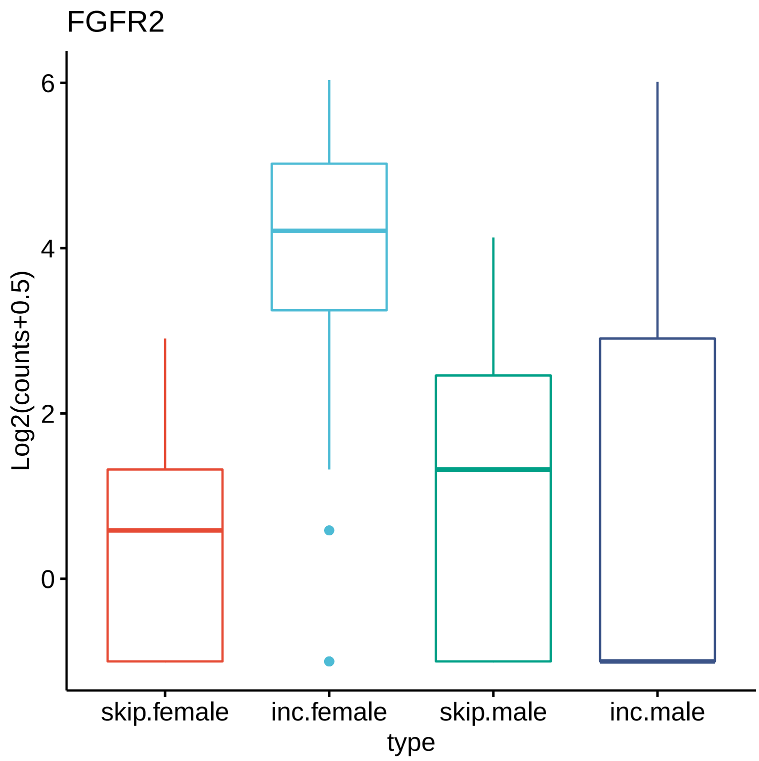


**Figure S19: FGFR2, IIIb** *FGFR2* skipped exon event in breast tissue (adjusted $p = 6.52 \times 10^{-15}$). This skipped exon event affects exon ENSE00003605080.1 (chr10:121518682-121518829). Five of the isoforms containing this exon are coding (ENST00000369059.5, ENST00000369058.7, ENST00000369056.5, ENST00000360144.7, ENST00000457416.6) and one is annotated as NMD ( ENST00000604236.5). The segment of the protein encoded by the exon is located C terminal to the tyrosine kinase domain of FGFR2. The exon encodes the sequence HSGINSSNAEVLALFNVTEADAGEYICKVSNYIGQANQSAWLTVLPKQQ which is represented in FGFR2 isoform 2 (NP_075259.4), 3 (NP_001138385.1), and 9 (NP_001138391.1). These correspond to the IIIb isoforms. Alternative splicing in FGFR2 generates the IIIb and IIIc isoforms. The IIIb isoforms are expressed exclusively in epithelial cells, while the IIIc isoforms are expressed only in mesenchymal cells. Mammary epithelial cells express FGFR2-IIIb, which binds FGF7 and FGF10, expressed by surrounding mesenchymal cells [14]. The location of the exon is shown in Figure S20.

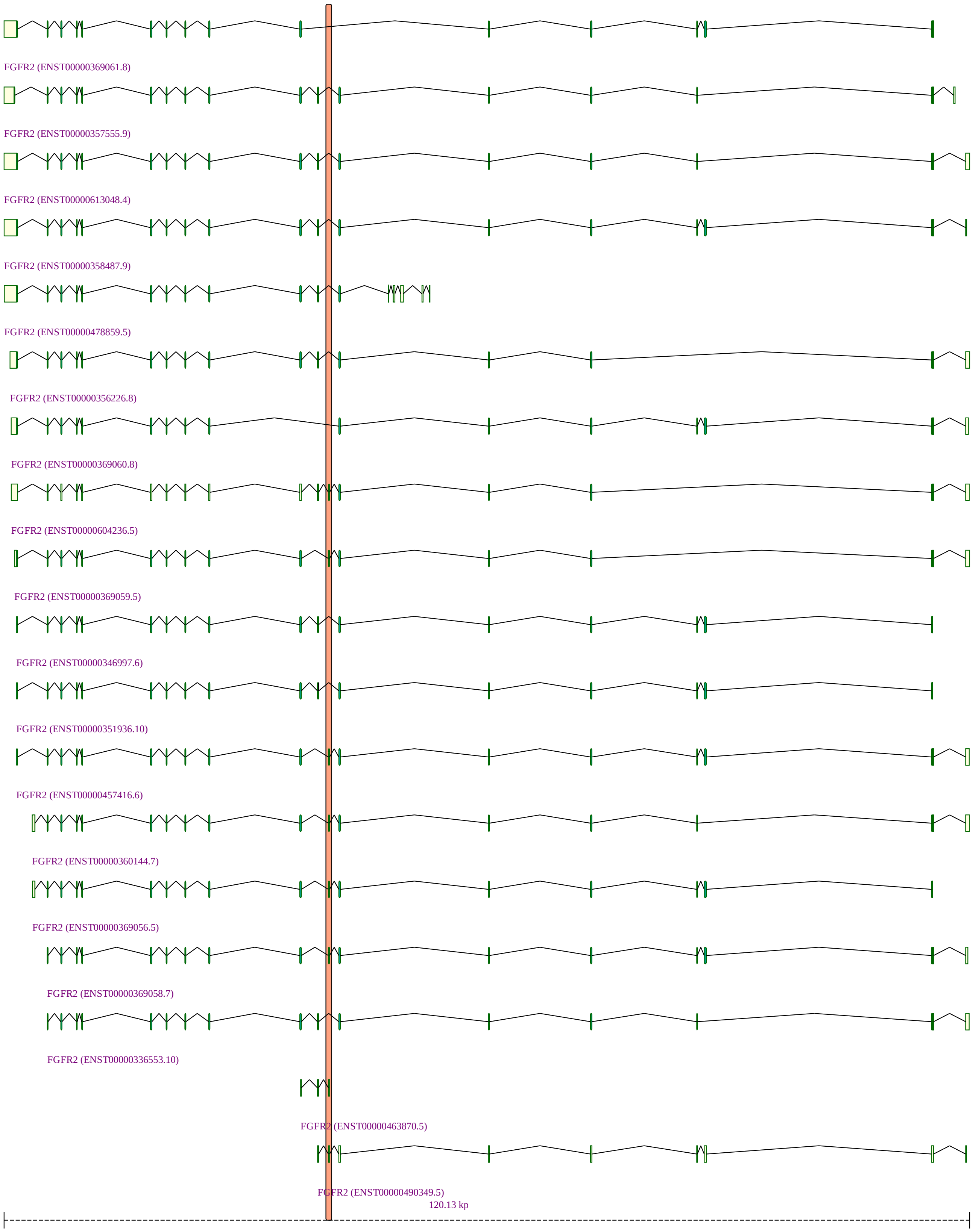


**Figure S20: FGFR2 (IIIb)**

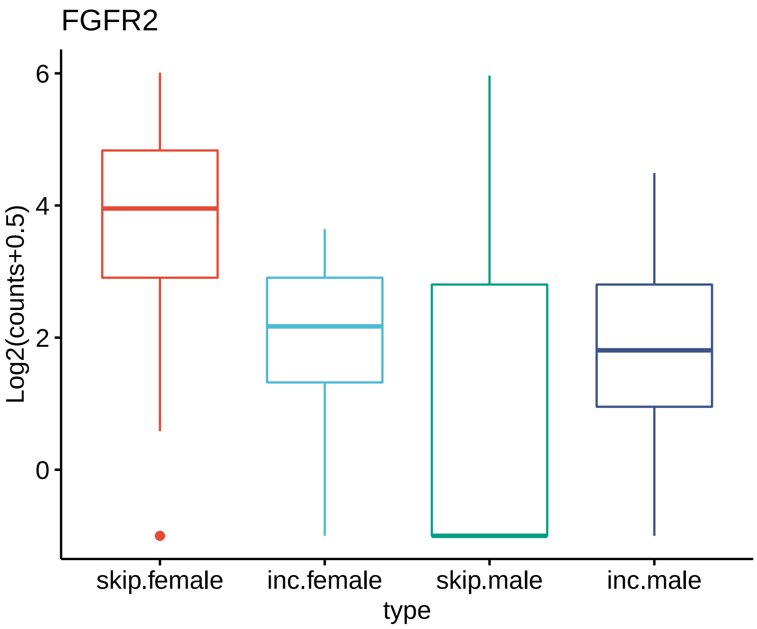


**Figure S21: FGFR2, IIIc** *FGFR2* skipped exon event in breast tissue (adjusted $p = 1.18 \times 10^{-10}$). This exon codes for IIIc isoforms of FGFR2, also known as FGFR2 (also called Bek/K-sam-I [15]). In breast tissue, females show a higher degree of skipping this exon, which corresponds to the higher degree of the exon associated with IIIb isoforms (See Figure S19. The location of the exon is shown in Figure S22.

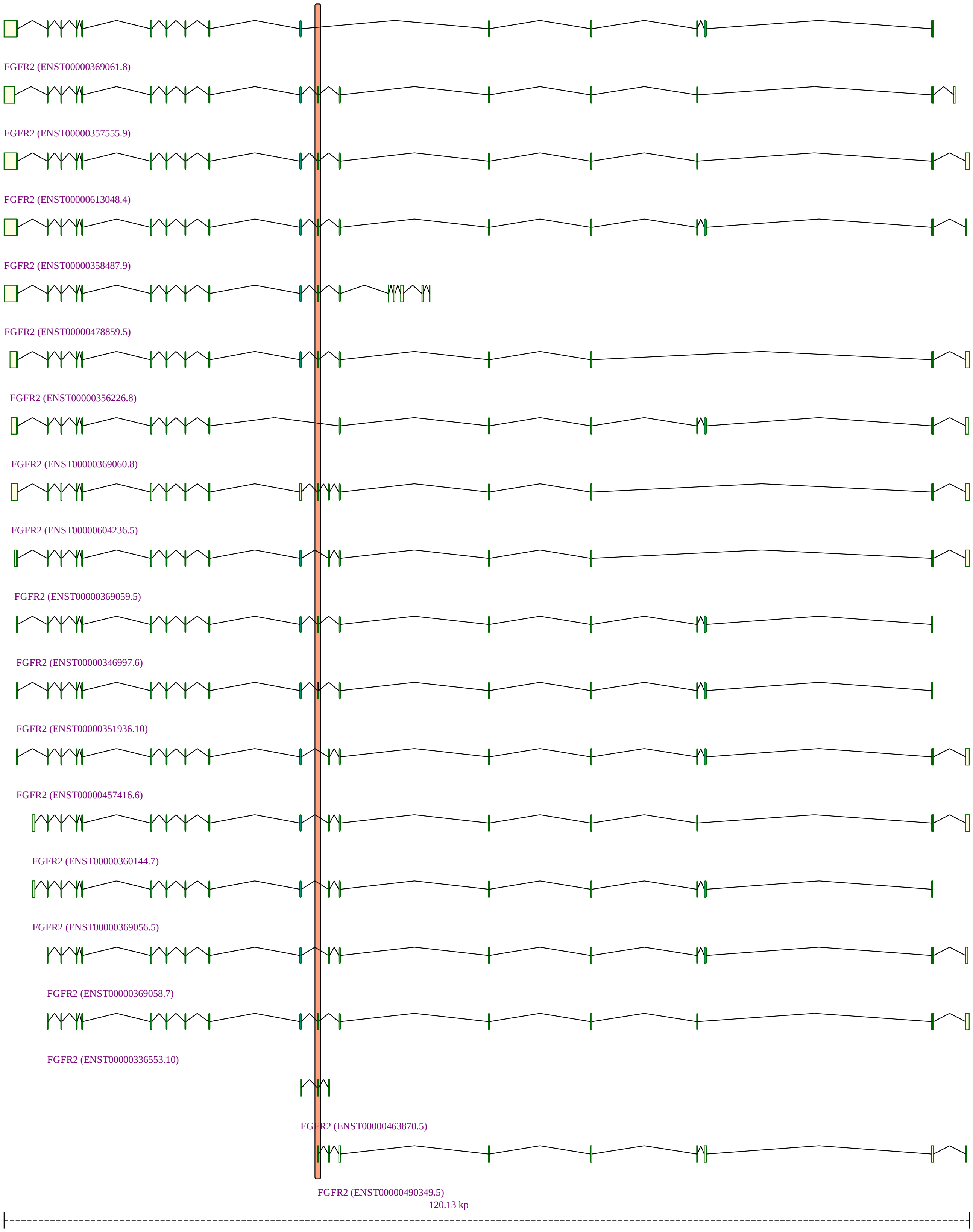


**Figure S22: FGFR2 (IIIc)**

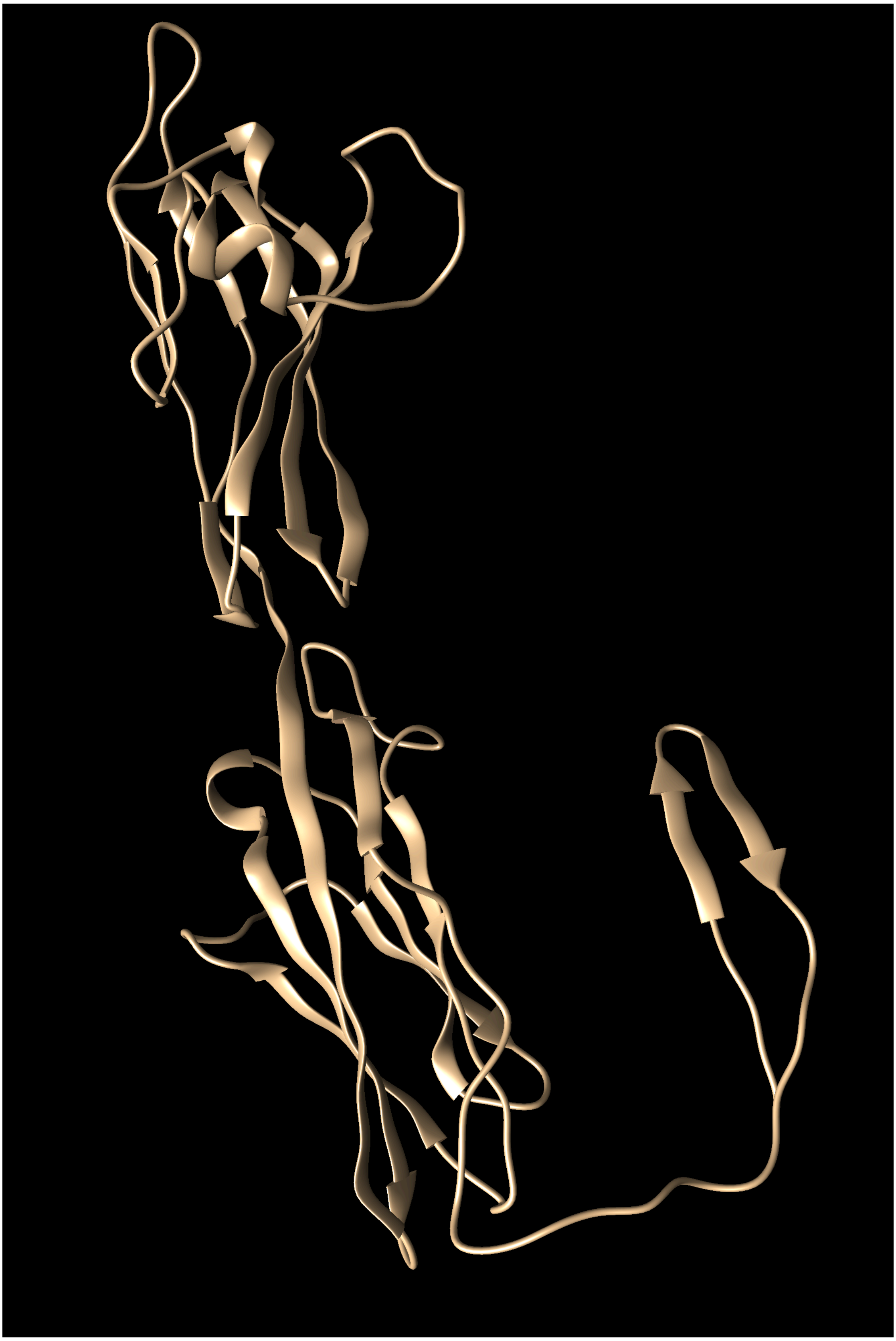

**Figure S23: FGFR2** Tertiary Structure of an FGFR2. The structure was obtained using homology modeling[7]. Graphic generated with the UCSF Chimera tool [6]

.

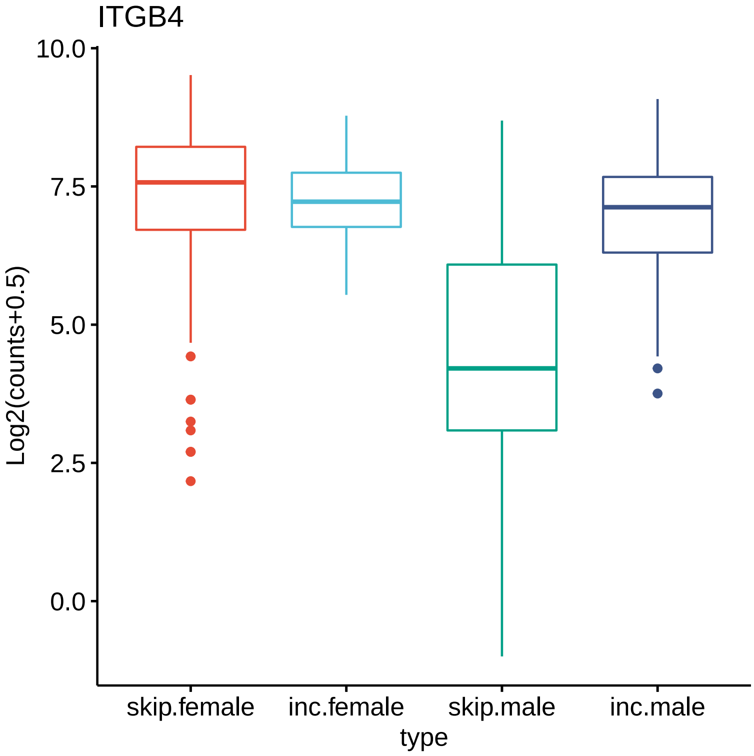


**Figure S24: ITGB4** Skipped exon event in *ITGB4* in breast tissue (adjusted $p = 5.63 \times 10^{-17}$). Integrins are transmembrane glycoprotein receptors that mediate cell-matrix or cell-cell adhesion and transduce signals that regulate gene expression and cell growth. These heterodimeric molecules consist of noncovalently linked alpha and beta subunits. Different combinations of alpha and beta polypeptides form complexes that vary in their ligand-binding specificities. Integrin $\alpha 6 \beta 4$ is one of the main laminin receptors and is primarily expressed by epithelial cells as an active component of hemidesmosomes [16]. The affected exon is ENSE00001385284 (chr17:75,755,05575,755,213). It is exon 33 of ENST00000449880.6. No information was available about the functional specifity of this isoform and the sequence encoded by this exon did not contain a motif as assessed by the ScanProsite tool [17]. Nonetheless, distinct isoforms of integrin $\alpha 6$ have different functions [18]. Figure S25 shows the location of the affected exon.

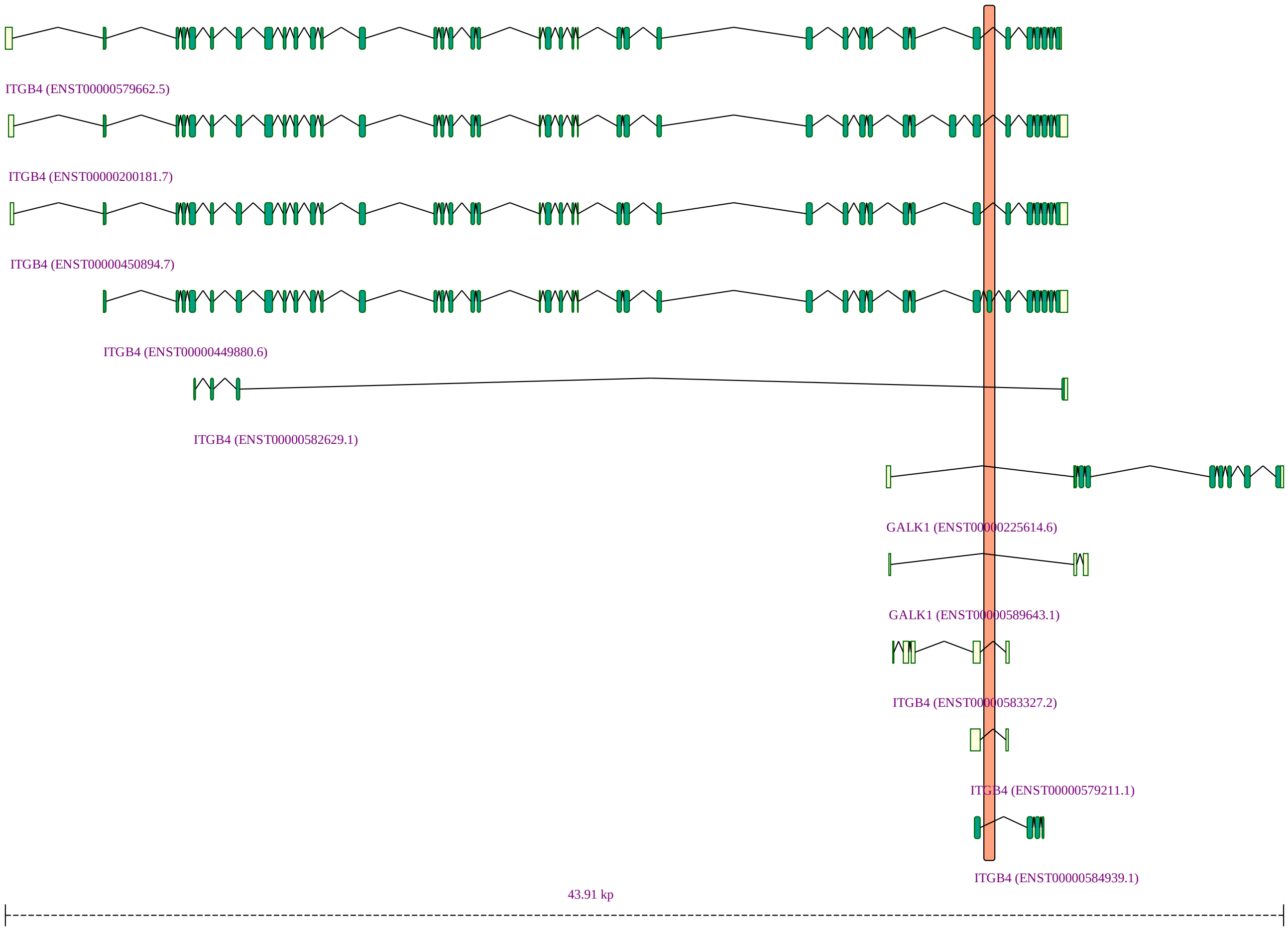


**Figure S25: ITGB4**

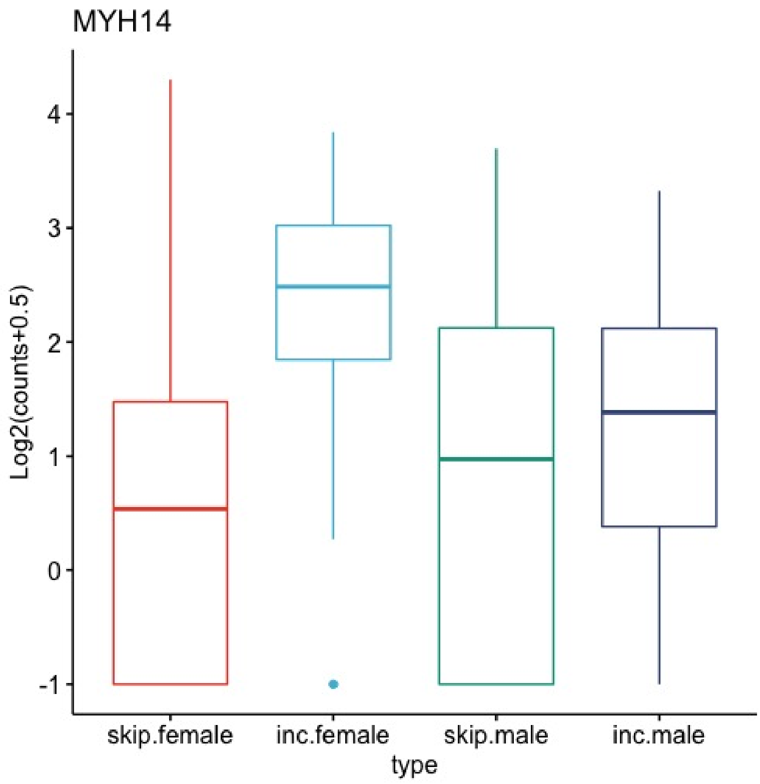


**Figure S26: MYH14** Skipped exon event in *MYH14* in breast tissue (adjusted $p = 3.12 \times 10^{-8}$). MYH14 is a member of the nonmuscle myosin II family of ATP-dependent molecular motors, which interact with cytoskeletal actin and regulate cytokinesis, cell motility, and cell polarity. The non muscle myosin heavy-chain gene *MYH14* has alternative spliced isoforms that differ by 8 amino acids located in the globular head of the protein. The affected exon, ENSE00001616058.1 (chr19:50224154-50224177), is 24 nucleotides in length and encodes these 8 amino acids. In myotonic dystrophy 1 (DM1), which is caused by a heterozygous trinucleotide repeat expansion (CTG)n in the 3'-UTR of |it DMPK, anomalous splicing of muscleblind-like 1 protein (MBNL1) and the CUG-binding protein 1 (CUGBP1); levels of the MBNL1 protein positively regulate the inclusion of this MYH14 exon, such that alterations of the MYH14 gene may contribute to the DM1 molecular pathogenesis [19]. The inserted exon is located near to Interestingly, this the isoform with exon inclusion is near to the ATP-binding region of MYH14 and increases both actin-activated (Mg2+)ATPase enzymatic activity and in vitro motility in translocating actin filaments [20]. In breast tissue, the ductal and acinar units are lined by two cell layers: the inner layer of epithelial cells lining the lumen and an outer layer of contractile myoepithelial cells (MEC). These cells contain myosin heavy chains [21], although to our knowledge, a specific role of MYH14 in mammary MECs has not been investigated to date. Our results suggest that female breast tissue is enriched for the inclusion isoforms that are associated with higher motility, which could support a role in lactation. Figure S27 shows the location of the affected exon.

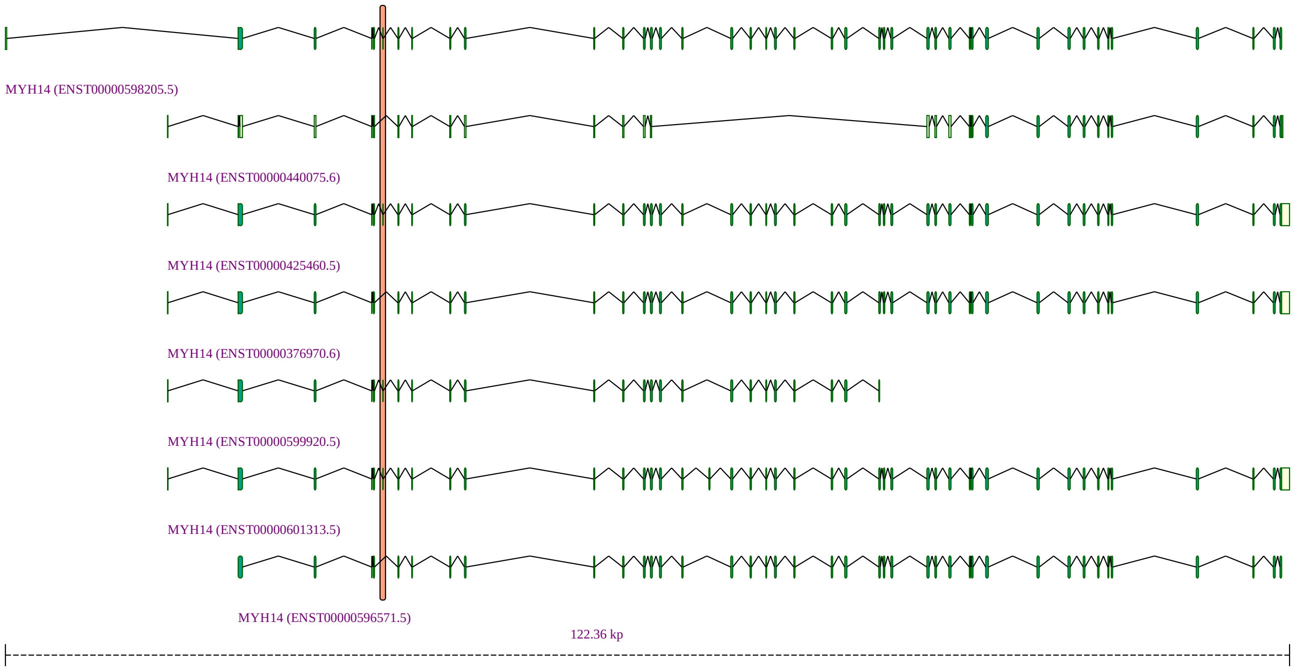


**Figure S27: MYH14**

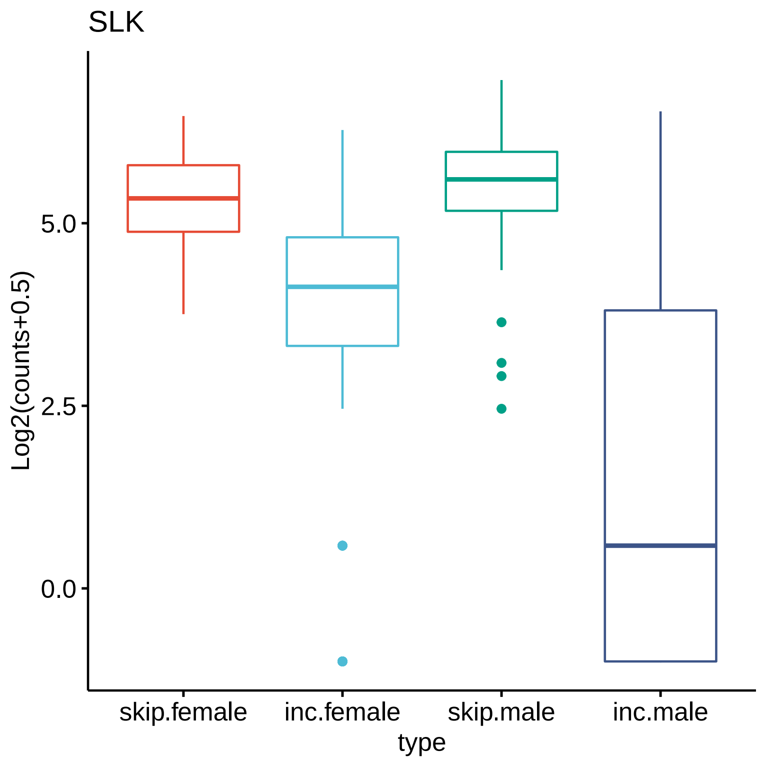


**Figure S28: SLK** *SLK* skipped exon event in breast tissue (adjusted $p = 2.32 \times 10^{-13}$). SLK is reported to be a mitogen-activated protein kinase kinase kinase kinase (MAP4K), and SLK can activate certain MAPK signaling cascades. SLK is predicted to regulate cytoskeletal organization and responses to apoptotic stimuli [22]. The affected exon is ENSE00000987678.1 (chr10:104010816-104010908), it is 93 nt in length. The exon inclusion levels of certain regulatory proteins can affect the activities of many transcription factors [23]. SRSF3 promotes skipping of this exon in cancer [24]. Inclusion of this exon was shown to promote nuclear over cytoplasmic localization (note that in the original publication, the exon is referred to as SLK_15_down, but it can be confirmed to be identical by comparison of the genomic coordinated in Supplemental table 4 (which are presented in hg19, rather than hg38 as in the present work). The location of the affected exon is presented in Figure S29.

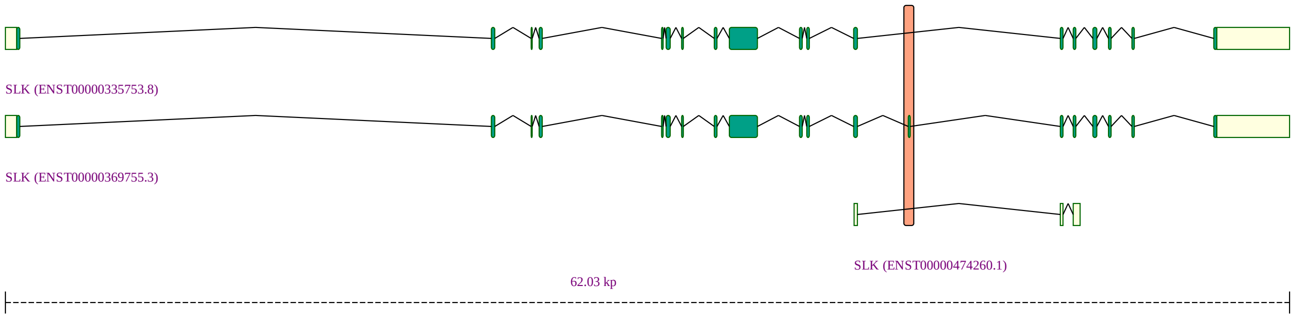


**Figure S29: SLK**

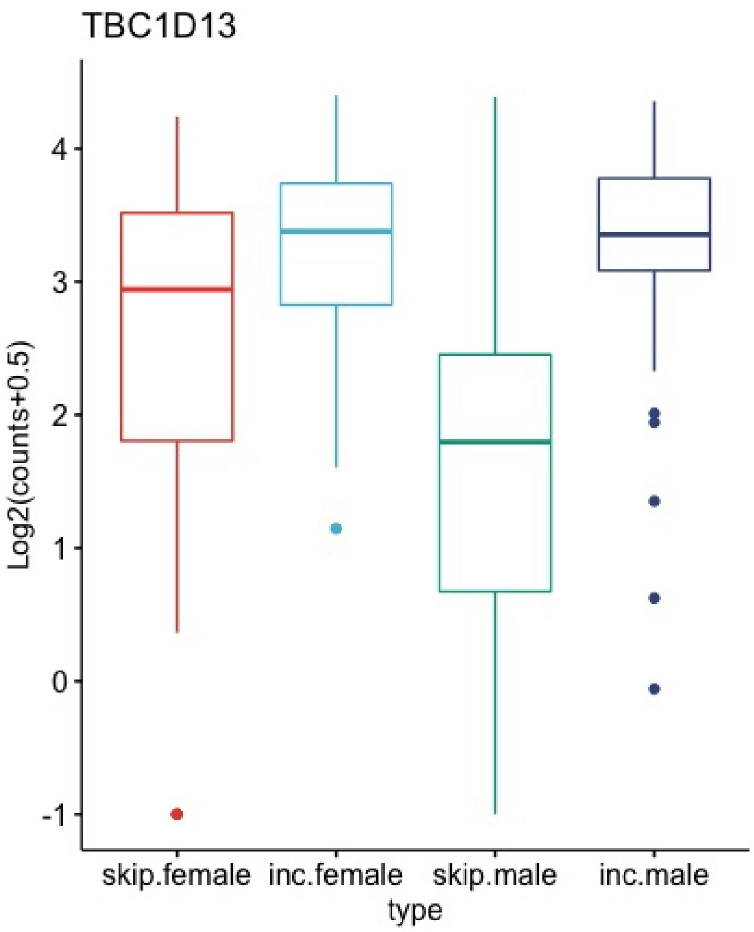


**Figure S30: TBC1D13** *TBC1D13* skipped exon event in breast tissue (adjusted $p = 1.19 \times 10^{-5}$) TBC1D13 is a Rab GTPase-activating protein (GAP) that stimulates GTP hydrolysis by Rab proteins. The skipped exon is at 128790734-128790775 (ENSE00003615055). The nonsense isoform ENST00000475097.5 is the only isoform that does not contain the exon. Thus, exon skipping is predicted to shift the distribution of TBC1D13 isoforms to non-protein coding forms. Tbc1d13 functions to inhibit insulin-stimulated Glut4 translocation to the plasma membrane [25]. Glucose is taken up by mammary epithelial cells through a passive, facilitative process, which is driven by the downward glucose concentration gradient across the plasma membrane. This process is mediated by facilitative glucose transporters (GLUTs), of which there are 14 known isoforms. Mammary glands mainly express GLUT1 and GLUT8 [26]. The shift towards non-protein coding isoforms of *TBC1D13* in female breast tissue may therefore suggest a physiological role of TBC1D13 and GLUT4 in breast tissue. The location of the affected exon is presented in Figure S31.

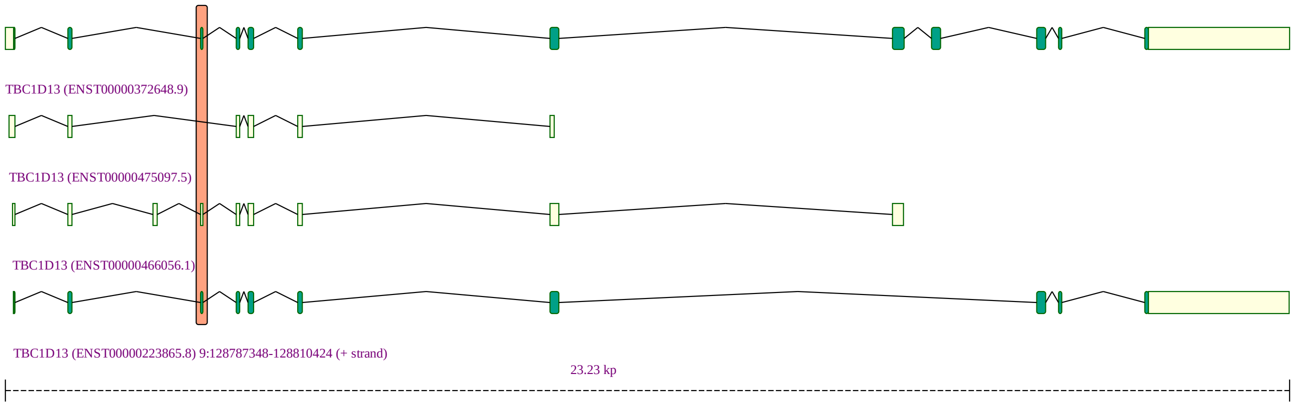


**Figure S31: TBC1D13**

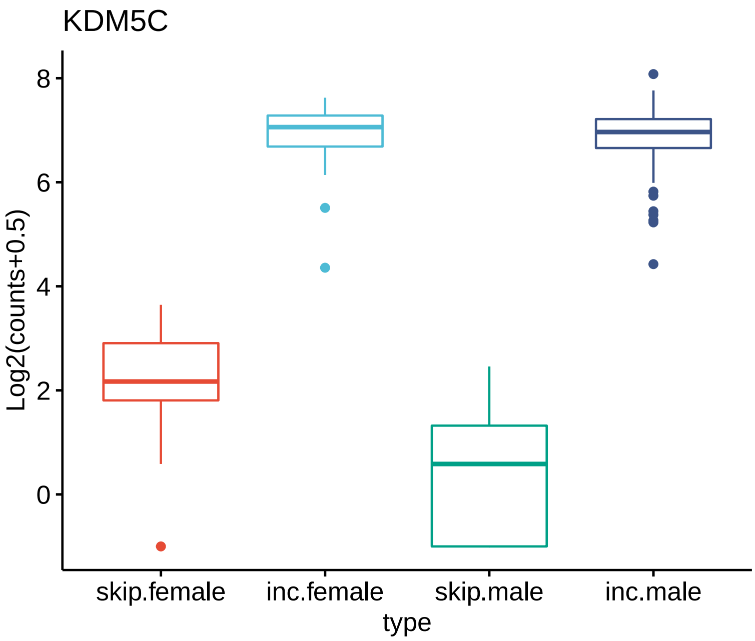


**Figure S32: KDM5C** *KDM5C* skipped exon event in cerebellum (adjusted $p = 1.95 \times 10^{-15}$; see Figure S32 for other tissues). KDM5C is a ubiquitously expressed X-chromosomal gene that encodes a specific H3K4me3 and H3K4me2 demethylase, and acts as a transcriptional repressor. The affected exon is ENSE00003572204.1 (chrX:53218276-53218398), and it is present in ENST00000467093.1 (non-protein coding processed transcripts) and ENST00000429877.5 (nonsense-mediated decay). We note there is a partially overlapping exon (ENSE00001637068 chrX:53,218,315-53,218,398 that is associated with the NMD transcript ENST00000428012.1). The proportion of skipped isoforms (not associated with NMD) is higher in females. This event was characterized as significant in multiple tissues (Figure S33). Figure S34 shows the location of the affected exon.

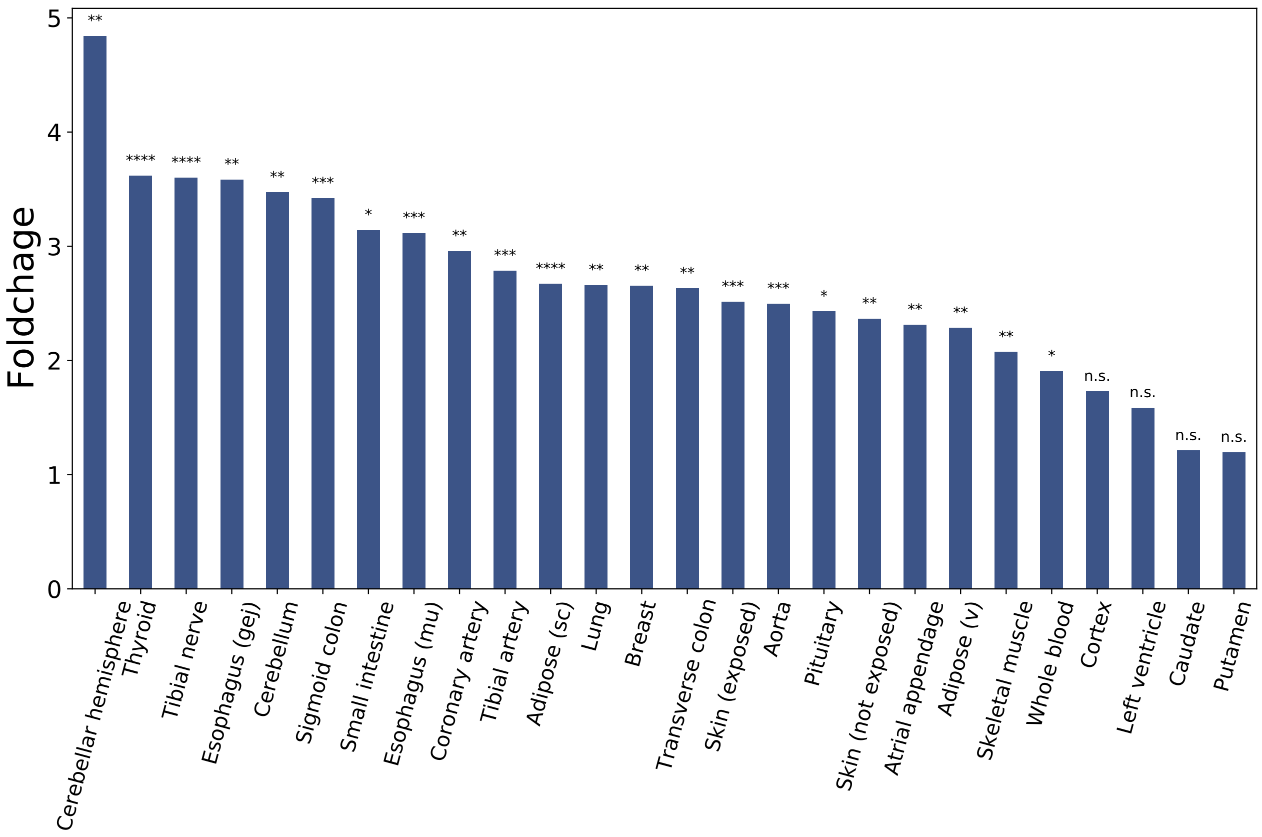


**Figure S33: KDM5C** This plot shows the average fold change of the *KDM5C* skipped exon event described in Figures S32 and S34. Statistical significance of the event is shown above the individual bars, with n.s.: not significant, *: $p < 10^{-5}$, **: $p < 10^{-10}$, ***: $p < 10^{-20}$, **** $p < 10^{-30}$ (adjusted).

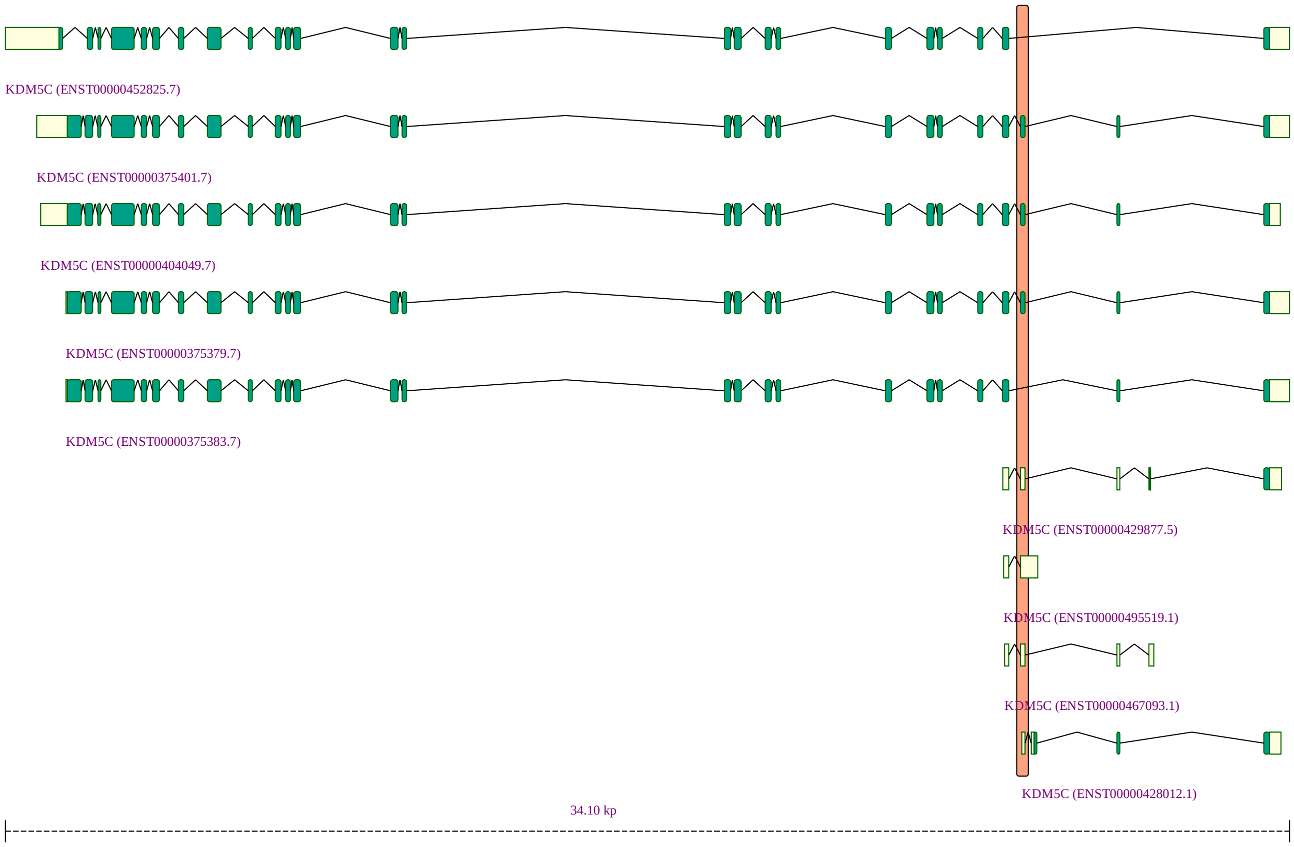


**Figure S34: KDM5C**

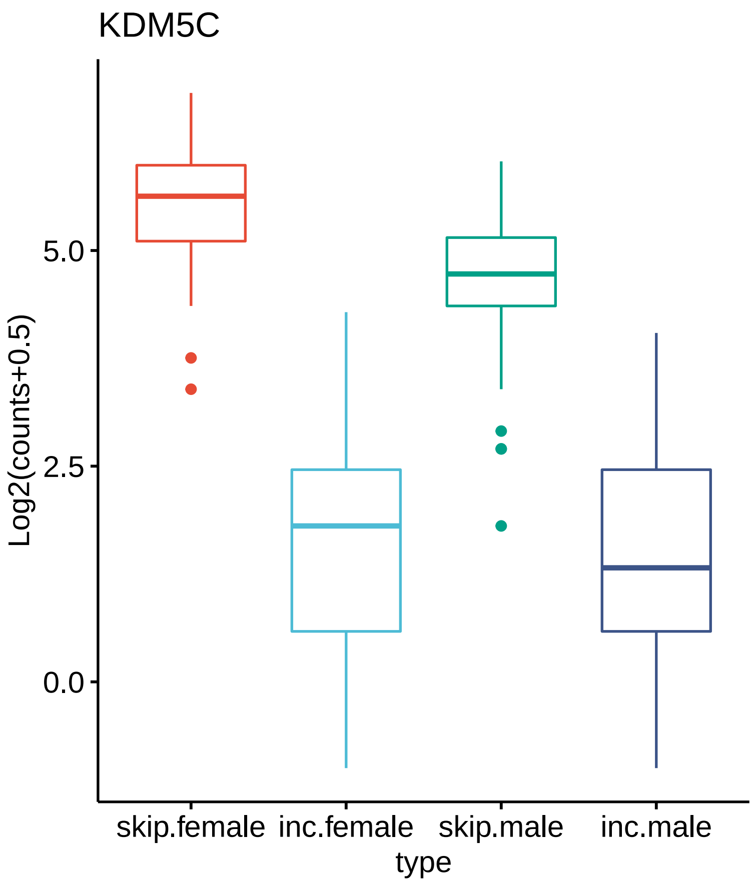


**Figure S35: KDM5C (Event 22850)** *KDM5C* skipped exon event in breast (adjusted $p$ = 0.0052). The affected exon is located at chrX:53221688-53221730. This exon is included in only one transcript, ENST00000429877.5, which is annotated as nonsense-mediated decay. Figure S36 shows the location of the affected exon. Similarly to the KDM5C event shown in Figures S34 and S32, females display a higher degree of nonskipped isoforms.

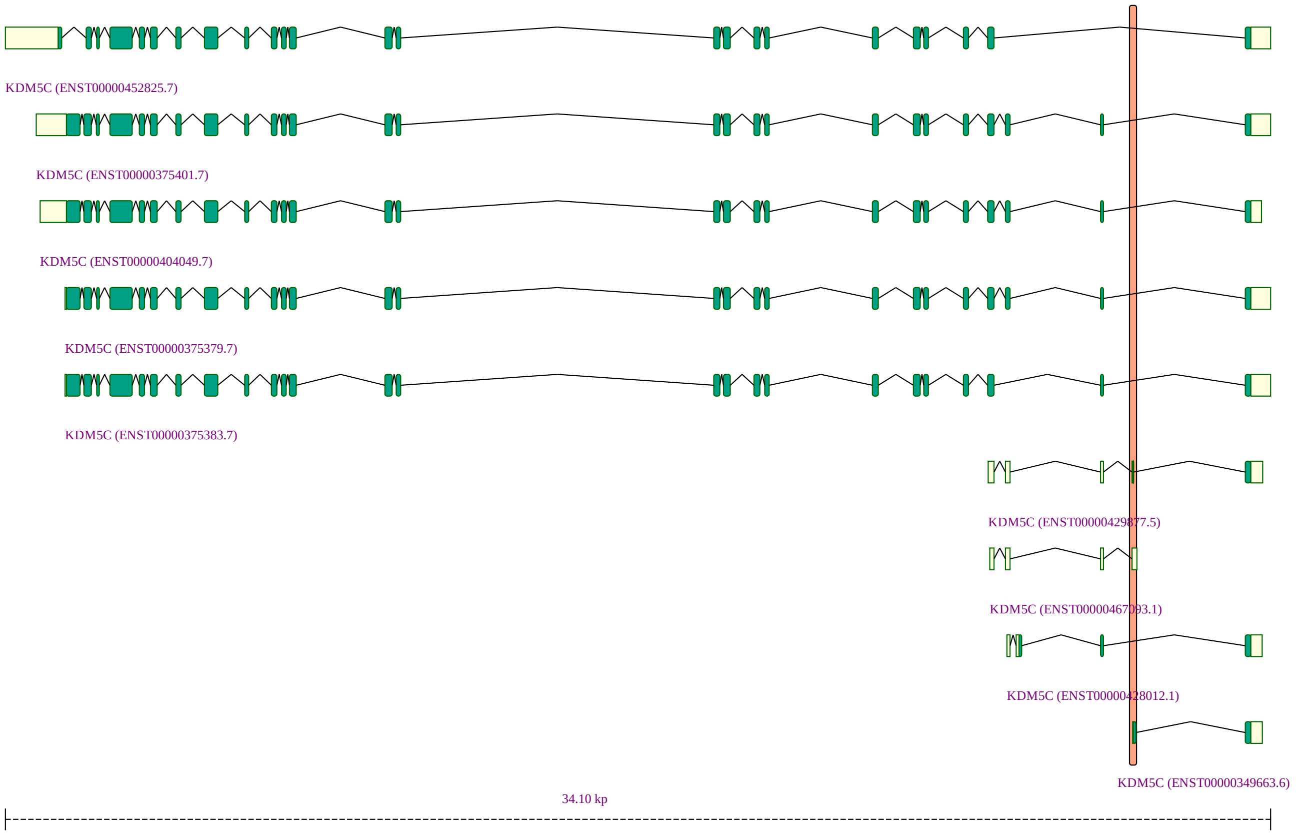


**Figure S36: KDM5C (Event 22850)**

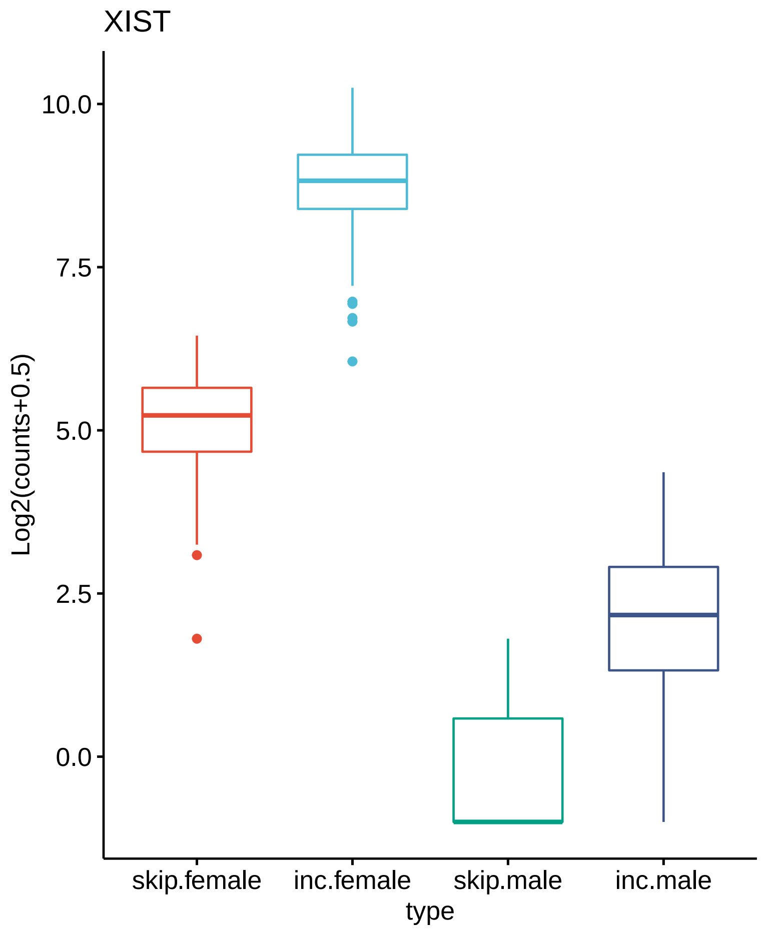


**Figure S37: XIST (Event 22850)** *XIST* A3SS event in esophagus (muscularis) (adjusted $p = 5.13 \times 10^{-13}$). The event was additionally found to be significant in Coronary artery (adjusted $p = 2.46 \times 10^{-17}$), Sigmoid colon(adjusted $p = 0.012$), Esophagus (gastroesophageal junction)(adjusted $p = 0.0021$), and atrial appendage (adjusted $p = 3.60 \times 10^{-22}$). THe event affects ENSE00003839041.1 (chrX:73826115-73827984).

| Gene | chr. | chromatin remodeling (GO:0006338) or covalent chromatin modification (GO:0016569) | regulation of gene expression (GO:0010468) |
|---|---|---|---|
| *XIST* | X | inactivation of X chromosome by DNA methylation (GO:0060821) | dosage compensation by inactivation of X chromosome (GO:0009048) |
| *KDM5C* | X | histone demethylase activity (GO:0032452) | negative regulation of transcription by RNA polymerase II (GO:0000122) |
| *DDX3X* | X | — | positive regulation of transcription by RNA polymerase II (GO:0045944) |
| *KDM6A* | X | chromatin remodeling (GO:0006338) | positive regulation of transcription by RNA polymerase II (GO:0045944) |
| *JPX* | X | inactivation of X chromosome by DNA methylation (GO:0060821) | regulation of dosage compensation by inactivation of X chromosome (GO:1900095) |

**Table S1:** Roles in signaling and gene regulation of genes with $> 10$ AS events. Representative annotations Gene Ontology (GO)are shown that are from the subhierarchies emanating from the GO terms chromatin remodeling (GO:0006338)/covalent chromatin modification (GO:0016569), signaling (GO:0023052), or regulation of gene expression (GO:0010468). 3 of the 9 genes had at least one annotation involving chromatic remodelling/modification, and 6 of the 9 genes had annotations related to gene expression regulation. 7 of the 9 genes were X chromosomal.

| Gene | n tissues | chromosome |
|---|---|---|
| *XIST* | 23 | chrX |
| *KDM5C* | 22 | chrX |
| *DDX3X* | 4 | chrX |
| *KDM6A* | 4 | chrX |
| *JPX* | 3 | chrX |
| *MYBPC1* | 2 | chr12 |
| *NAT9* | 2 | chr17 |
| *SEPT6* | 2 | chrX |
| *FLII* | 2 | chr17 |
| *NDRG4* | 2 | chr16 |
| *PIP5KL1* | 2 | chr9 |
| *RAB26* | 2 | chr16 |
| *ALG13* | 2 | chrX |
| *CACNB1* | 2 | chr17 |
| *DEPDC5* | 2 | chr22 |
| *DPM1* | 2 | chr20 |
| *ENAH* | 2 | chr1 |
| *GCOM1* | 2 | chr15 |
| *KMT2E* | 2 | chr7 |
| *MPDU1* | 2 | chr17 |
| *PTPN6* | 2 | chr12 |

**Table S2:** Genes displaying significant alternative splicing in 2 or more tissues. 7 of these 21 gene are encoded by the X chromosome.

| Tissue | Males | Females | Included | Display |
|---|---|---|---|---|
| Adipose-Subcutaneous | 218 | 445 | 1 | Adipose (sc) |
| Adipose-Visceral(Omentum) | 170 | 371 | 1 | Adipose (v) |
| AdrenalGland | 101 | 157 | 1 | Adrenal gland |
| Artery-Aorta | 153 | 279 | 1 | Aorta |
| Artery-Coronary | 94 | 146 | 1 | Coronary artery |
| Artery-Tibial | 209 | 454 | 1 | Tibial artery |
| Bladder | 7 | 14 | 0 | n/a |
| Brain-Amygdala | 45 | 107 | 0 | n/a |
| Brain-Anteriorcingulatecortex(BA24) | 48 | 128 | 0 | n/a |
| Brain-Caudate(basalganglia) | 63 | 183 | 1 | Caudate |
| Brain-CerebellarHemisphere | 58 | 157 | 1 | Cerebellar hemisphere |
| Brain-Cerebellum | 67 | 174 | 1 | Cerebellum |
| Brain-Cortex | 74 | 181 | 1 | Cortex |
| Brain-FrontalCortex(BA9) | 56 | 153 | 1 | Frontal cortex |
| Brain-Hippocampus | 54 | 143 | 1 | Hippocampus |
| Brain-Hypothalamus | 55 | 147 | 1 | Hypothalamus |
| Brain-Nucleusaccumbens(basalganglia) | 64 | 182 | 1 | Nucleus accumbens |
| Brain-Putamen(basalganglia) | 49 | 156 | 1 | Putamen |
| Brain-Spinalcord(cervicalc-1) | 57 | 102 | 1 | Spinal cord |
| Brain-Substantianigra | 38 | 101 | 0 | n/a |
| Breast-MammaryTissue | 168 | 291 | 1 | Breast |
| Cells-Culturedfibroblasts | 174 | 330 | 1 | Fibroblasts |
| Cells-EBV-transformedlymphocytes | 62 | 112 | 1 | EBV-lymphocytes |
| Colon-Sigmoid | 133 | 240 | 1 | Sigmoid colon |
| Colon-Transverse | 147 | 259 | 1 | Transverse colon |
| Esophagus-GastroesophagealJunction | 124 | 251 | 1 | Esophagus (gej) |
| Esophagus-Mucosa | 192 | 363 | 1 | Esophagus (m) |
| Esophagus-Muscularis | 177 | 338 | 1 | Esophagus (mu) |
| Heart-AtrialAppendage | 136 | 293 | 1 | Atrial appendage |
| Heart-LeftVentricle | 138 | 294 | 1 | Left ventricle |
| Kidney-Cortex | 19 | 66 | 0 | n/a |
| Kidney-Medulla | 1 | 3 | 0 | n/a |
| Liver | 65 | 161 | 1 | Liver |
| Lung | 183 | 395 | 1 | Lung |
| MinorSalivaryGland | 47 | 115 | 0 | n/a |
| Muscle-Skeletal | 260 | 543 | 1 | Skeletal muscle |
| Nerve-Tibial | 200 | 419 | 1 | Tibial nerve |
| Ovary | 180 | 0 | 0 | n/a |
| Pancreas | 121 | 207 | 1 | Pancreas |
| Pituitary | 79 | 204 | 1 | Pituitary |
| Prostate | 0 | 245 | 0 | n/a |
| Skin-NotSunExposed(Suprapubic) | 193 | 411 | 1 | Skin (not exposed) |
| Skin-SunExposed(Lowerleg) | 234 | 467 | 1 | Skin (exposed) |
| SmallIntestine-TerminalIleum | 67 | 120 | 1 | Small intestine |
| Spleen | 87 | 154 | 1 | Spleen |
| Stomach | 132 | 227 | 1 | Stomach |
| Testis | 0 | 361 | 0 | n/a |
| Thyroid | 219 | 434 | 1 | Thyroid |
| Uterus | 142 | 0 | 0 | n/a |
| WholeBlood | 254 | 501 | 1 | Whole blood |

**Table S3:** GTEx samples used in this project. The columns `Male` and `Female` indicate the number of avaiable samples. Tissues were analyzed if at least 50 samples were available for both sexes (indicated by a “1” in the `Included` column; a “0” indicates that a tissue was not included. The `Display name` column indicates the possibly shortened name used to refer to the tissue throughout the text and in the figures.

| Tissue | positive regulation of gonadotropin secretion | extracellular matrix | eukaryotic translation initiation factor 2 complex | septin cytoskeleton | heparan sulfate sulfotransferase activity | septin ring | regulation of follicle-stimulating hormone secretion | collagen-containing extracellular matrix | positive regulation of follicle-stimulating hormone secretion | septin complex | regulation of gonadotropin secretion | multicellular organism development | developmental process | anatomical structure development | negative regulation of B cell differentiation |
|---|---|---|---|---|---|---|---|---|---|---|---|---|---|---|---|
| tibial nerve | | ✓ | ✓ | | ✓ | | | ✓ | | | | ✓ | ✓ | ✓ | |
| fibroblasts | | | ✓ | | | | | | | | | ✓ | ✓ | ✓ | |
| spinal cord | | | | | | | | | | | | | | | |
| Small intestine | | | | | | | | | | | | | | | |
| Nucleus accumbens | | | | | | | | | | | | | | | |
| Esophagus (mu) | ✓ | | | ✓ | | ✓ | ✓ | | ✓ | ✓ | ✓ | | | | ✓ |
| Adipose (v) | | | | | ✓ | | | | | | | | | | |
| Esophagus (m) | | | | | | | | | | | | | | | |
| Hypothalamus | | | | | | | | | | | | | | | |
| Pancreas | | | | | ✓ | | | | | | | | | | |
| Atrial appendage | | ✓ | | | ✓ | | | ✓ | | | | ✓ | ✓ | ✓ | |
| Cerebellum | | | | | | | | | | | | | | | |
| Left ventricle | ✓ | ✓ | ✓ | | | | ✓ | ✓ | ✓ | | ✓ | | | | ✓ |
| Pituitary | | ✓ | | | | | | ✓ | | | | ✓ | ✓ | ✓ | ✓ |
| Transverse colon | | | ✓ | | | | | | | | | | | | |
| Adipose (sc) | ✓ | ✓ | ✓ | ✓ | ✓ | ✓ | ✓ | | ✓ | ✓ | ✓ | ✓ | ✓ | ✓ | ✓ |
| Frontal cortex | | ✓ | | | | | | ✓ | | | | | | | |
| Esophagus (gej) | | | | ✓ | | ✓ | | | | ✓ | | | | | |
| Spleen | | | | | | | | | | | | | | | |
| Liver | | | | | | | | | | | | | | | |
| EBV-lymphocytes | | | ✓ | | | | | | | | | | | | |
| Tibial artery | ✓ | ✓ | ✓ | ✓ | | ✓ | ✓ | ✓ | ✓ | ✓ | ✓ | ✓ | ✓ | ✓ | ✓ |
| Breast | ✓ | ✓ | ✓ | ✓ | ✓ | ✓ | ✓ | ✓ | ✓ | ✓ | ✓ | ✓ | ✓ | ✓ | ✓ |
| Skeletal muscle | ✓ | | | | ✓ | | ✓ | | ✓ | | ✓ | | | | |
| Skin (exposed) | ✓ | ✓ | ✓ | ✓ | ✓ | ✓ | ✓ | ✓ | ✓ | ✓ | ✓ | ✓ | ✓ | ✓ | ✓ |
| Whole blood | ✓ | | | ✓ | ✓ | ✓ | ✓ | | ✓ | ✓ | ✓ | | | | |
| Lung | | | ✓ | | | | | | | | | | | | |
| Thyroid | | | | ✓ | ✓ | ✓ | | | | ✓ | ✓ | | | | ✓ |
| Aorta | | ✓ | | | | | | ✓ | | | | | | | |
| Cerebellar hemisphere | | | | | | | | | | | | | | | |
| Skin (not exposed) | ✓ | ✓ | ✓ | ✓ | ✓ | ✓ | ✓ | | ✓ | ✓ | ✓ | ✓ | ✓ | ✓ | ✓ |
| Hippocampus | | | | | | | | | | | | | | | |

**Table S4:** Summary of Gene Ontology analysis for differential gene expression. A blue cell with a checkmark indicates that the indicated GO term was significantly enriched in the tissue shown by the row in question. Terms that were significant in the highest number of tissues are shown.

**Table S5:** GO Analysis of differentially spliced genes.

| GO term | tissue | study | population | p-value | adj. p-value |
|---|---|---|---|---|---|
| posttranscriptional regulation of gene expression GO:0010608 | Esophagus (mu) | 18/138(13.0%) | 433/11270(3.8%) | 5.0E-6 | 0.009013 |
| membrane-bounded organelle GO:0043227 | Esophagus (mu) | 121/138(87.7%) | 8239/11270(73.1%) | 2.4E-5 | 0.040154 |
| intracellular membrane-bounded organelle GO:0043231 | Esophagus (mu) | 113/138(81.9%) | 7308/11270(64.8%) | 7.0E-6 | 0.012165 |
| RNA binding GO:0003723 | Esophagus (mu) | 33/138(23.9%) | 1195/11270(10.6%) | 5.0E-6 | 0.008801 |
| nucleus GO:0005634 | Esophagus (mu) | 84/138(60.9%) | 4841/11270(43.0%) | 1.5E-5 | 0.025873 |
| protein demethylase activity GO:0140457 | Atrial appendage | 2/5(40.0%) | 23/11270(0.2%) | 4.0E-5 | 0.004604 |
| histone methyltransferase complex GO:0035097 | Atrial appendage | 2/5(40.0%) | 58/11270(0.5%) | 2.58E-4 | 0.029897 |
| histone demethylation GO:0016577 | Atrial appendage | 2/5(40.0%) | 25/11270(0.2%) | 4.7E-5 | 0.005458 |
| histone lysine demethylation GO:0070076 | Atrial appendage | 2/5(40.0%) | 23/11270(0.2%) | 4.0E-5 | 0.004604 |
| protein demethylation GO:0006482 | Atrial appendage | 2/5(40.0%) | 27/11270(0.2%) | 5.5E-5 | 0.006383 |
| dioxygenase activity GO:0051213 | Atrial appendage | 2/5(40.0%) | 65/11270(0.6%) | 3.24E-4 | 0.037573 |
| protein dealkylation GO:0008214 | Atrial appendage | 2/5(40.0%) | 27/11270(0.2%) | 5.5E-5 | 0.006383 |
| demethylase activity GO:0032451 | Atrial appendage | 2/5(40.0%) | 30/11270(0.3%) | 6.8E-5 | 0.007907 |
| histone demethylase activity GO:0032452 | Atrial appendage | 2/5(40.0%) | 23/11270(0.2%) | 4.0E-5 | 0.004604 |
| demethylation GO:0070988 | Atrial appendage | 2/5(40.0%) | 49/11270(0.4%) | 1.84E-4 | 0.021304 |
| protein-containing complex GO:0032991 | Spleen | 36/64(56.3%) | 3561/11270(31.6%) | 3.8E-5 | 0.042163 |
| macromolecule metabolic process GO:0043170 | Spleen | 44/64(68.8%) | 4194/11270(37.2%) | 0.0 | 3.2E-4 |
| organic substance metabolic process GO:0071704 | Spleen | 47/64(73.4%) | 5346/11270(47.4%) | 2.0E-5 | 0.02272 |
| response to steroid hormone GO:0048545 | Spleen | 7/64(10.9%) | 165/11270(1.5%) | 3.9E-5 | 0.043767 |
| nitrogen compound metabolic process GO:0006807 | Spleen | 44/64(68.8%) | 4715/11270(41.8%) | 1.2E-5 | 0.013208 |
| metabolic process GO:0008152 | Spleen | 50/64(78.1%) | 5718/11270(50.7%) | 6.0E-6 | 0.006388 |
| gene expression GO:0010467 | Spleen | 22/64(34.4%) | 1448/11270(12.8%) | 8.0E-6 | 0.008731 |
| cellular metabolic process GO:0044237 | Spleen | 48/64(75.0%) | 5206/11270(46.2%) | 2.0E-6 | 0.002771 |
| primary metabolic process GO:0044238 | Spleen | 45/64(70.3%) | 5076/11270(45.0%) | 3.7E-5 | 0.041108 |
| cellular macromolecule metabolic process GO:0044260 | Spleen | 36/64(56.3%) | 3421/11270(30.4%) | 1.4E-5 | 0.015922 |
| NF-kappaB binding GO:0051059 | Spleen | 4/64(6.3%) | 26/11270(0.2%) | 1.3E-5 | 0.014405 |
| RNA destabilization GO:0050779 | EBV-lymphocytes | 2/5(40.0%) | 29/11270(0.3%) | 6.4E-5 | 0.006554 |
| protein-containing complex GO:0032991 | Breast | 472/1253(37.7%) | 3561/11270(31.6%) | 1.0E-6 | 0.004626 |
| cellular component morphogenesis GO:0032989 | Breast | 74/1253(5.9%) | 394/11270(3.5%) | 3.0E-6 | 0.020738 |
| cell-substrate junction GO:0030055 | Breast | 69/1253(5.5%) | 341/11270(3.0%) | 0.0 | 0.002785 |
| enzyme binding GO:0019899 | Breast | 234/1253(18.7%) | 1616/11270(14.3%) | 4.0E-6 | 0.024855 |
| anchoring junction GO:0070161 | Breast | 111/1253(8.9%) | 607/11270(5.4%) | 0.0 | 3.05E-4 |
| GTPase binding GO:0051020 | Breast | 79/1253(6.3%) | 422/11270(3.7%) | 2.0E-6 | 0.011026 |
| focal adhesion GO:0005925 | Breast | 67/1253(5.3%) | 335/11270(3.0%) | 1.0E-6 | 0.006339 |
| cadherin binding GO:0045296 | Breast | 59/1253(4.7%) | 258/11270(2.3%) | 0.0 | 2.4E-4 |
| nucleic acid metabolic process GO:0090304 | Skin (exposed) | 10/20(50.0%) | 1541/11270(13.7%) | 1.11E-4 | 0.043661 |
| cellular macromolecule biosynthetic process GO:0034645 | Skin (exposed) | 9/20(45.0%) | 1084/11270(9.6%) | 4.3E-5 | 0.016923 |
| gene expression | | | | | |

*Continued on next page*

Table S5 – *Continued from previous page*

| GO term | tissue | study | population | p-value | adj. p-value |
|---|---|---|---|---|---|
| GO:0010467 macromolecule biosynthetic process | Skin (exposed) | 10/20(50.0%) | 1448/11270(12.8%) | 6.5E-5 | 0.025441 |
| GO:0009059 RNA metabolic process | Skin (exposed) | 9/20(45.0%) | 1108/11270(9.8%) | 5.1E-5 | 0.020158 |
| GO:0016070 mRNA metabolic process | Skin (exposed) | 9/20(45.0%) | 1092/11270(9.7%) | 4.6E-5 | 0.017948 |
| GO:0016071 response to lithium ion | Skin (exposed) | 7/20(35.0%) | 527/11270(4.7%) | 2.1E-5 | 0.008426 |
| GO:0010226 negative regulation of protein acetylation | Thyroid | 2/17(11.8%) | 10/11270(0.1%) | 9.6E-5 | 0.042106 |
| GO:1901984 negative regulation of peptidyl-lysine acetylation | Aorta | 2/12(16.7%) | 16/11270(0.1%) | 1.24E-4 | 0.033768 |
| GO:2000757 | Aorta | 2/12(16.7%) | 14/11270(0.1%) | 9.4E-5 | 0.025638 |

**Table S6:** Genes with three or more sex-biased alternative splicing events

| Gene | Tissue | AS | Position | IC (m/f) | SC (m/f) |
|---|---|---|---|---|---|
| KDM5C | skin not sun exposed suprapubic | SE | chrX:53218275-53218398 | 103.0/134.1 | 0.6/2.8 |
| KDM5C | thyroid | SE | chrX:53218275-53218398 | 106.3/148.1 | 0.3/3.5 |
| KDM5C | artery aorta | SE | chrX:53218275-53218398 | 85.1/117.0 | 0.7/3.6 |
| KDM5C | breast mammary tissue | SE | chrX:53218275-53218398 | 78.0/129.3 | 0.4/3.2 |
| KDM5C | breast mammary tissue | SE | chrX:53221687-53221730 | 3.2/3.9 | 27.9/51.5 |
| KDM5C | adipose visceral omentum | SE | chrX:53218275-53218398 | 71.2/102.1 | 0.3/2.5 |
| KDM5C | muscle skeletal | SE | chrX:53218275-53218398 | 113.5/145.1 | 0.8/2.8 |
| KDM5C | skin sun exposed lower leg | SE | chrX:53218275-53218398 | 104.9/143.3 | 0.4/2.7 |
| KDM5C | brain cerebellum | SE | chrX:53218275-53218398 | 124.1/126.5 | 1.3/5.1 |
| KDM5C | artery tibial | SE | chrX:53218275-53218398 | 91.8/125.1 | 0.8/3.8 |
| KDM5C | artery coronary | SE | chrX:53218275-53218398 | 81.0/106.0 | 0.5/3.3 |
| KDM5C | esophagus muscularis | SE | chrX:53218275-53218398 | 88.1/120.5 | 0.4/3.4 |
| KDM5C | whole blood | SE | chrX:53218275-53218398 | 55.6/68.3 | 0.6/2.5 |
| KDM5C | small intestine terminal ileum | SE | chrX:53218275-53218398 | 94.1/143.1 | 0.4/3.7 |
| KDM5C | heart atrial appendage | SE | chrX:53218275-53218398 | 56.9/75.5 | 0.3/2.2 |
| KDM5C | brain cerebellar hemisphere | SE | chrX:53218275-53218398 | 125.9/146.7 | 0.7/6.6 |
| KDM5C | lung | SE | chrX:53218275-53218398 | 97.9/137.2 | 0.8/3.8 |
| KDM5C | colon sigmoid | SE | chrX:53218275-53218398 | 76.7/111.2 | 0.4/3.6 |
| KDM5C | nerve tibial | SE | chrX:53218275-53218398 | 90.2/114.8 | 0.6/4.1 |
| KDM5C | pituitary | SE | chrX:53218275-53218398 | 100.7/133.4 | 0.2/2.1 |

Continued on next page

Table S6 – continued from previous page

| Gene | Tissue | Category | Position | IC (m/f) | SC (m/f) |
|---|---|---|---|---|---|
| KDM5C | adipose subcutaneous | SE | chrX:53218275-53218398 | 77.2/106.5 | 0.4/2.8 |
| KDM5C | esophagus gastroesophageal junction | SE | chrX:53218275-53218398 | 82.4/109.8 | 0.2/3.2 |
| KDM5C | colon transverse | SE | chrX:53218275-53218398 | 81.3/119.3 | 0.3/2.8 |
| XIST | thyroid | MXE | chrX:73831065-73831274 | 0.4/351.8 | 1.2/764.0 |
| XIST | artery aorta | MXE | chrX:73831065-73831274 | 0.4/193.9 | 0.8/415.6 |
| XIST | liver | MXE | chrX:73831065-73831274 | 0.3/98.6 | 0.5/227.2 |
| XIST | muscle skeletal | MXE | chrX:73831065-73831274 | 0.3/120.6 | 0.6/246.1 |
| XIST | spleen | MXE | chrX:73831065-73831274 | 0.7/124.3 | 1.2/288.9 |
| XIST | artery tibial | MXE | chrX:73831065-73831274 | 0.3/175.1 | 0.7/364.1 |
| XIST | brain putamen basal ganglia | MXE | chrX:73831065-73831274 | 0.3/67.1 | 0.4/143.2 |
| XIST | artery coronary | A3SS | chrX:73826114-73827984 | 1.7/483.5 | 0.2/52.0 |
| XIST | brain frontal cortex ba 9 | MXE | chrX:73831065-73831274 | 0.2/74.0 | 0.4/161.4 |
| XIST | esophagus muscularis | SE | chrX:73822070-73822216 | 1.1/137.8 | 0.3/47.5 |
| XIST | esophagus muscularis | A3SS | chrX:73826114-73827984 | 5.0/471.6 | 0.4/38.2 |
| XIST | pancreas | MXE | chrX:73831065-73831274 | 0.3/137.6 | 0.6/295.7 |
| XIST | brain caudate basal ganglia | MXE | chrX:73831065-73831274 | 0.3/93.6 | 0.6/206.2 |
| XIST | whole blood | MXE | chrX:73831065-73831274 | 0.2/21.3 | 0.4/47.4 |
| XIST | heart atrial appendage | A3SS | chrX:73826114-73827984 | 3.1/443.2 | 0.3/39.9 |
| XIST | brain cerebellar hemisphere | MXE | chrX:73831065-73831274 | 0.2/139.1 | 0.5/365.4 |
| XIST | lung | MXE | chrX:73831065-73831274 | 0.4/186.9 | 0.6/417.6 |
| XIST | colon sigmoid | SE | chrX:73833237-73833374 | 4.4/400.6 | 0.2/32.8 |
| XIST | colon sigmoid | A3SS | chrX:73826114-73827984 | 6.7/519.5 | 0.5/46.7 |
| XIST | colon sigmoid | A3SS | chrX:73831065-73831274 | 5.7/438.2 | 0.2/12.0 |
| XIST | cells cultured fibroblasts | MXE | chrX:73831065-73831274 | 0.2/190.4 | 0.5/431.5 |
| XIST | esophagus mucosa | MXE | chrX:73831065-73831274 | 0.3/141.2 | 0.9/325.2 |
| XIST | nerve tibial | MXE | chrX:73831065-73831274 | 7.2/275.0 | 21.8/558.9 |
| XIST | nerve tibial | A3SS | chrX:73831065-73831274 | 25.5/637.8 | 0.5/15.7 |
| XIST | brain hippocampus | MXE | chrX:73831065-73831274 | 0.3/69.7 | 0.5/158.4 |
| XIST | pituitary | MXE | chrX:73831065-73831274 | 0.3/420.4 | 0.7/864.9 |

Continued on next page

**Table S6 – continued from previous page**

| Gene | Tissue | Category | Position | IC (m/f) | SC (m/f) |
|---|---|---|---|---|---|
| XIST | esophagus gastroesophageal junction | A3SS | chrX:73826114-73827984 | 5.2/457.3 | 0.4/40.9 |
| DDX3X | breast mammary tissue | RI | chrX:41346228-41346622 | 8.9/7.7 | 202.0/263.1 |
| DDX3X | breast mammary tissue | A3SS | chrX:41346493-41346622 | 0.7/0.4 | 202.0/263.1 |
| DDX3X | muscle skeletal | SE | chrX:41339035-41339083 | 183.3/252.0 | 0.5/3.3 |
| DDX3X | muscle skeletal | SE | chrX:41339038-41339083 | 79.4/110.5 | 0.5/3.3 |
| DDX3X | muscle skeletal | SE | chrX:41337690-41339083 | 85.4/119.1 | 0.5/3.3 |
| DDX3X | muscle skeletal | A5SS | chrX:41337407-41339083 | 127.0/176.5 | 0.5/3.3 |
| DDX3X | esophagus muscularis | RI | chrX:41346228-41346622 | 4.0/3.8 | 168.0/254.3 |
| DDX3X | cells cultured fibroblasts | RI | chrX:41343215-41344128 | 4.2/6.7 | 0.3/0.4 |
| KDM6A | liver | SE | chrX:45060021-45060156 | 2.7/6.5 | 0.6/0.8 |
| KDM6A | muscle skeletal | A5SS | chrX:45090722-45090864 | 22.2/40.4 | 0.4/0.7 |
| KDM6A | brain caudate basal ganglia | A5SS | chrX:45090722-45090864 | 15.7/27.1 | 0.5/0.4 |
| KDM6A | heart atrial appendage | A5SS | chrX:45090722-45090864 | 18.1/28.4 | 0.5/0.6 |
| JPX | artery tibial | SE | chrX:73994839-73994961 | 13.8/24.3 | 0.6/0.8 |
| JPX | lung | SE | chrX:73994839-73994961 | 14.3/22.7 | 0.5/0.5 |
| JPX | brain nucleus accumbens basal ganglia | SE | chrX:73994839-73994961 | 22.5/46.7 | 0.5/0.5 |

| **Gene** | **ensembl gene id** | **ensembl exon id** | **position** | **AS** |
|---|---|---|---|---|
| *DDX3X* | ENSG00000215301.10 | ENSE00003458274.1 | chrX:41339036-41339083 | SE |
| **affected transcripts** | | • protein coding: n=17<br>• lncRNA: n=3<br>• nonsense-mediated decay: n=12<br>• retained intron: n=8 | | |
| *DDX3X* | ENSG00000215301.10 | ENSE00003458274.1 | chrX:41339036-41339083 | SE |
| **affected transcripts** | | • protein coding: n=17<br>• lncRNA: n=3<br>• nonsense-mediated decay: n=12<br>• retained intron: n=8 | | |
| *DDX3X* | ENSG00000215301.10 | ENSE00003822339.1 | chrX:41337691-41339083 | SE |
| **affected transcripts** | | • nonsense-mediated decay: n=1<br>• retained intron: n=1 | | |
| *DDX3X* | ENSG00000215301.10 | ENSE00003825146.1 | chrX:41337408-41339083 | A5SS |
| **affected transcripts** | | • retained intron: n=1 | | |
| *DDX3X* | ENSG00000215301.10 | ENSE00003820599.1 | chrX:41343216-41344128 | RI |
| **affected transcripts** | | • retained intron: n=1 | | |
| *DDX3X* | ENSG00000215301.10 | ENSE00003830261.1 | chrX:41346494-41346622 | RI |
| **affected transcripts** | | • nonsense-mediated decay:: n=1 | | |
| *DDX3X* | ENSG00000215301.10 | ENSE00003824510.1 | chrX:41346229-41346622 | RI |
| **affected transcripts** | | • retained intron: n=1 | | |

**Table S7:** Sex-biased alternative splicing involving *DDX3X*. In Ensembl, *DDX3X* has 23 protein-coding, 3 lncRNA, 14 nonsense-mediated decay, and 29 retained intron transcripts. The significant alternative splicing events listed here were identified in one tissue each except for , which was identified in mammary tissue and esophagus (muscularis).

| Gene | ensembl gene id | ensembl exon id | position | AS |
|---|---|---|---|---|
| *XIST* | ENSG00000229807.12 | ENSE00001742809.1 | chrX:73831066-73831274 | MXE |
| **affected transcripts** | | • lncRNA: n=16 | | |
| *XIST* | ENSG00000229807.12 | ENSE00003831669.1 | chrX:73826114-73827984 | A3SS |
| **affected transcripts** | | • lncRNA: n=9 | | |
| *XIST* | ENSG00000229807.12 | ENSE00001641042.1: | chrX:73822070-73822216 | SE |
| **affected transcripts** | | • lncRNA: n=5 | | |
| *XIST* | ENSG00000229807.12 | ENSE00001692247.1 | chrX:73833237-73833374 | SE |
| **affected transcripts** | | • lncRNA: n=15 | | |

**Table S8:** Sex-biased alternative splicing involving *XIST*. Four different significant AS events were detected. The event affecting ENSE00001742809.1 was identified in 20 tissues, the event affecting ENSE00001641042.1 was only found in esophagus (muscularis), the event affecting ENSE00003831669.1 was identified in 5 tissues, and the event affecting ENSE00001692247.1 was only found in sigmoid colon.

| Cell Line | DAST GTEx | Overlap DAST (GTEx/Meng) | DAST Meng | Total Genes Meng | p-value |
|---|---|---|---|---|---|
| N88 | 574 | 72 | 602 | 7071 | 0.0003911611 |
| N182 572 | 100 | 1017 | 572 | 7056 | 0.01879892 |
| N129 | 549 | 129 | 1357 | 6725 | 0.02606251 |

**Table S9:** We analyzed published RNA-seq data from samples from ER+ cells in cultures established from primary cells in which cells were exposed to estrogen to assess estrogen-induced gene expression changes for a total of three cell lines [27]. We refer to this dataset according to the name of the first author as Meng. Isoform read counts were generated with RSEM and differential gene expression and differential splicing were assessed with HBA-DEALS [28]. We defined the DAST group to be genes for which differential splicing but not differential expression was identified by HBA-DEALS. Overlap was assessed by Fisher's exact test, and a $p$-value for the set of three experiments was obtained by Fisher's combined probability test.

| Id | Symbol | Gene |
|---|---|---|
| 7164 | TPD52L1 | TPD52 like 1 |
| 3983 | ABLIM1 | actin binding LIM protein 1 |
| 89797 | NAV2 | neuron navigator 2 |
| 120 | ADD3 adducin 3 | |
| 2317 | FLNB | filamin B |
| 9612 | NCOR2 | nuclear receptor corepressor 2 |
| 960 | CD44 | CD44 molecule |
| 771 | CA12 | carbonic anhydrase 12 |
| 4137 | MAPT | microtubule associated protein tau |
| 51195 | RAPGEFL1 | Rap guanine nucleotide exchange factor like 1 |
| 6337 | SCNN1A | sodium channel epithelial 1 subunit alpha |
| 4582 | MUC1 | mucin 1, cell surface associated |
| 1999 | ELF3 | E74 like ETS transcription factor 3 |
| 3866 | KRT15 | keratin 15 |
| 64284 | RAB17 | RAB17, member RAS oncogene family |
| 79083 | MLPH | melanophilin |
| 80004 ESRP2 | epithelial splicing regulatory protein 2 | |

**Table S10:** MSigDB Pathway analysis of genes differentially spliced in breast tissue revealed HALLMARK_ESTROGEN_RESPONSE_EARLY (early) as the top enriched pathway, with a total of 17 associated genes.

| |
|---|
| https://cloudos.lifebit.ai/app/jobs/5e79013586fdca01036e78f0 |
| https://cloudos.lifebit.ai/app/jobs/5e79158c86fdca01036e99d7 |
| https://cloudos.lifebit.ai/app/jobs/5e79dcb886fdca01036f98ef |
| https://cloudos.lifebit.ai/app/jobs/5e7a5e66acc18a010404e673 |
| https://cloudos.lifebit.ai/app/jobs/5e7b875aacc18a0104067279 |
| https://cloudos.lifebit.ai/app/jobs/5e7b8780acc18a01040672a1 |
| https://cloudos.lifebit.ai/app/jobs/5e7bbe62acc18a010406d547 |
| https://cloudos.lifebit.ai/app/jobs/5e7bc0a4acc18a010406daa3 |
| https://cloudos.lifebit.ai/app/jobs/5e7bc0cdacc18a010406db31 |
| https://cloudos.lifebit.ai/app/jobs/5e7cd41bacc18a010409f1a6 |
| https://cloudos.lifebit.ai/app/jobs/5e7d03beacc18a01040a5b5d |
| https://cloudos.lifebit.ai/app/jobs/5e7d25d2acc18a01040ab1b5 |
| https://cloudos.lifebit.ai/app/jobs/5e8191e7c1a31e1bf69b03dc |
| https://cloudos.lifebit.ai/app/jobs/5e8191d89dfe0b1bf3b148b0 |
| https://cloudos.lifebit.ai/app/jobs/5e8191caef806a1bef41c706 |
| https://cloudos.lifebit.ai/app/jobs/5e7f7a32e6162a091cc7f47b |
| https://cloudos.lifebit.ai/app/jobs/5e7f7c38c9fc200944ef49ae |
| https://cloudos.lifebit.ai/app/jobs/5e7f7c64bedf9c0948586f8f |
| https://cloudos.lifebit.ai/app/jobs/5e7f7d742ef52d09587343b2 |

**Table S11:** Links to Nextflow analyses performed as batches of up to 500 samples on the Lifebit CloudOS platform

[7] Andrew Waterhouse, Martino Bertoni, Stefan Bienert, Gabriel Studer, Gerardo Tauriello, Rafal Gumienny, Florian T Heer, Tjaart A P de Beer, Christine Rempfer, Lorenza Bordoli, Rosalba Lepore, and Torsten Schwede. SWISS-MODEL: homology modelling of protein structures and complexes. *Nucleic Acids Research*, 46(W1):W296–W303, May 2018.

[8] E C Dell'Angelica, H Ohno, C E Ooi, E Rabinovich, K W Roche, and J S Bonifacino. Ap-3: an adaptor-like protein complex with ubiquitous expression. *The EMBO journal*, 16:917–928, March 1997.

[9] Guus Martinus Bol, Min Xie, and Venu Raman. Ddx3, a potential target for cancer treatment. *Molecular cancer*, 14:188, November 2015.

[10] Xia Wang, Jennifer E Posey, Jill A Rosenfeld, Carlos A Bacino, Fernando Scaglia, LaDonna Immken, Jill M Harris, Scott E Hickey, Theresa M Mosher, Anne Slavotinek, Jing Zhang, Joke Beuten, Magalie S Leduc, Weimin He, Francesco Vetrini, Magdalena A Walkiewicz, Weimin Bi, Rui Xiao, Pengfei Liu, Yunru Shao, Alper Gezdirici, Elif Y Gulec, Yunyun Jiang, Sandra A Darilek, Adam W Hansen, Michael M Khayat, Davut Pehlivan, Juliette Piard, Donna M Muzny, Neil Hanchard, John W Belmont, Lionel Van Maldergem, Richard A Gibbs, Mohammad K Eldomery, Zeynep C Akdemir, Adekunle M Adesina, Shan Chen, Yi-Chien Lee, Undiagnosed Diseases Network, Brendan Lee, James R Lupski, Christine M Eng, Fan Xia, Yaping Yang, Brett H Graham, and Paolo Moretti. Phenotypic expansion in ddx3x - a common cause of intellectual disability in females. *Annals of clinical and translational neurology*, 5:1277–1285, October 2018.

[16] Jean-François Beaulieu. Integrin $\alpha6\beta4$ in colorectal cancer: Expression, regulation, functional alterations and use as a biomarker. *Cancers*, 12, December 2019.
